# Regulatory variants active in iPSC-derived pancreatic progenitor cells are associated with Type 2 Diabetes in adults

**DOI:** 10.1101/2021.10.20.465206

**Authors:** Jennifer P. Nguyen, Agnieszka D’Antonio-Chronowska, Kyohei Fujita, Bianca M. Salgado, Hiroko Matsui, Timothy D. Arthur, iPSCORE Consortium, Margaret K.R. Donovan, Matteo D’Antonio, Kelly A. Frazer

## Abstract

Pancreatic progenitor cells (PPC) are an early developmental multipotent cell type that give rise to mature endocrine, exocrine, and ductal cells. To investigate the extent to which regulatory variants active in PPC contribute to pancreatic complex traits and disease in the adult, we derived PPC from induced pluripotent stem cells (iPSCs) of nine unrelated individuals and generated single cell profiles of chromatin accessibility (snATAC- seq) and transcriptome (scRNA-seq). While iPSC-PPC differentiation was asynchronous and included cell types from early to late developmental stages, we found that the predominant cell type consisted of NKX6-1+ progenitors. Genetic characterization using snATAC-seq identified 86,261 regulatory variants that either displayed chromatin allelic bias and/or were predicted to affect active transcription factor (TF) binding sites. Integration of these regulatory variants with 380 fine-mapped type 2 diabetes (T2D) risk loci identified regulatory variants in 209 of these loci that are functional in iPSC-PPC, either by affecting transcription factor binding or through association with allelic effects on chromatin accessibility. The PPC active regulatory variants in 65 of these loci showed strong evidence of causally underlying the association with T2D. Our study shows that studying the functional associations of regulatory variation in iPSC-PPC enables the identification and characterization of causal SNPs for adult Type 2 Diabetes.

## Introduction

In early development the pancreas is formed from pancreatic progenitor cells (PPCs), which are a multipotent cell type that has the potential to give rise to endocrine cells (clusters of hormone secreting cells, such α, β, γ, and δ cells) and exocrine cells (i.e. acinar and ductal cells) (Cano et al., 2014; Jennings et al., 2015). While PPCs are precursors to mature pancreas cell types, the extent to which regulatory variants active in PPCs contribute to pancreatic complex traits and disease in the adult is currently not known. Recently it has become possible to use human pluripotent stem cells to derive PPCs (Pagliuca et al., 2014; Rezania et al., 2014), which provide a virtually unlimited source of cells to identify and characterize regulatory variants. Given that regulatory variation is largely located in enhancers and promoters (Pennacchio et al., 2013), ATAC-seq provides an optimal method to identify and characterize variants in PPCs that directly alter transcription factor binding and downstream gene expression. Examining induced pluripotent stem cell derived PPCs (iPSC-PPCs) from whole-genome sequenced unrelated individuals using ATAC-seq could enable identification of regulatory variation in PPCs and determine whether or not they are associated with adult pancreatic traits and diseases such as Type 2 Diabetes (T2D).

PPCs are characterized as a population of cells that have differentiated beyond the pancreatic foregut, committed to a pancreatic progenitor fate, and marked by the co-expression of *PDX1* and *NKX6-1* (Cano et al., 2014; Jennings et al., 2015). A reference set of embryonic stem cell-derived PPCs (ESC-PPC) obtained across multiple differentiation stages have shown the presence of multiple cell types, including pancreatic progenitors, endocrine cells and exocrine cells (Veres et al., 2019), suggesting that stem cell differentiation of PPCs is likely asynchronous. It is currently unclear how closely iPSC-PPC will be to ESC-PPC, i.e., whether similar cell types resulting from asynchronous differentiation will be observed. Furthermore, the reproducibility of pancreatic differentiation across iPSC lines derived from different individuals is unknown.

The development of single nuclear ATAC-seq (snATAC-seq) has become a powerful tool to investigate the mechanisms underlying the function of regulatory variants (Chiou et al., 2021; Rai et al., 2020). snATAC-seq pinpoints the locations of regulatory elements across the genome; and integration with motif footprinting and transcription factor binding prediction tools enable the identification of transcription factor binding sites associated with each snATAC-seq peak as well as a determination of the allelic effects of each SNP on transcription factor binding (Bentsen et al., 2020; Ghandi et al., 2014; Ghandi et al., 2016; Yan et al., 2021). Therefore, snATAC-seq provides an optimal approach to characterize regulatory genetic variation and to identify molecular mechanisms (i.e. transcription factor binding) underlying the associations between genotype and T2D.

The genetic variants associated with the most common pancreatic disease, T2D, have been investigated in millions of people, resulting in the identification of more than 500 loci (Mahajan et al., 2018; Vujkovic et al., 2020). Although several studies have successfully identified the likely causal variant in a small subset of T2D- associated loci (Chiou et al., 2021; Mahajan et al., 2018; Varshney et al., 2017), the vast majority of the signals still remains uncharacterized and typically lie within regulatory regions. Given the importance of many transcription factors in regulating pancreatic cells’ function, it is not surprising that many non-coding validated T2D risk variants overlap transcription factor binding sites (Chiou et al., 2021; Geusz et al., 2020; Greenwald et al., 2019; Mahajan et al., 2018; Rai et al., 2020; Thurner et al., 2018), indicating that snATAC-seq provides an optimal method to identify the molecular mechanisms underlying the role of regulatory variants in this disease.

Here, we investigated the potential of iPSC-PPC as a model system to study the associations between genetic variation, gene regulation and T2D. We used nine iPSC lines from unrelated individuals (Panopoulos et al., 2017) and a 15-day differentiation protocol to obtain ten iPSC-PPC samples. We characterized these iPSC-PPC samples using scRNA-seq and snATAC-seq and found that, while differentiation occurs asynchronously across iPSC lines, the vast majority of derived cells are NKX6-1+ progenitors, which represent early pancreatic lineage to endocrine cells. We investigated the associations between genetic variation and the function of chromatin accessible regions in iPSC-PPC and observed that 86,261 regulatory variants either overlapped footprints of transcription factors active in iPSC-PPC, were predicted to have allelic effects on transcription factor binding sites, and/or were demonstrated to have allelic-specific effects on snATAC-seq signals. We next investigated the overlap between these variants and SNPs included in 380 functional T2D credible sets from a recent GWAS (Mahajan et al., 2018) and found that 209 had SNPs that either overlapped binding sites of active transcription factors and/or were associated with allelic effects, including 65 that showed strong evidence of being causal. This study demonstrates that many T2D risk variants overlap regulatory elements active in iPSC-PPC and display allelic effects due to altered transcription factor binding, indicating that iPSC-PPC are a suitable model system to study the genetics of T2D in adults.

## Results

We used iPSC lines from nine unrelated iPSCORE individuals (Panopoulos et al., 2017) (Table S1) and a 15-day differentiation protocol to derive ten iPSC-derived pancreatic progenitor (iPSC-PPC) samples (one iPSC clone was differentiated twice; Figure 1A). To perform a baseline assessment of iPSC-PPC differentiation efficiency, we measured the fraction of cells positive for two hallmark PPC transcription factors, PDX1 and NKX6-1, using flow cytometry, and observed that the ten samples differentiated with varying efficiency (Figure S1). The fraction of cells that stained positive for PDX1 ranged from 22.1 to 96.4%, and the fraction that was double-positive stained for PDX1 and NKX6-1 ranged from 9.4 to 91.7% (Figure S1B). As expression of NKX6-1 occurs later than PDX1 expression during development, the lower fraction of double-positive cells reflect asynchronous differentiation resulting in pancreatic progenitor cells at slightly different stages of maturity.

**Figure 1:**
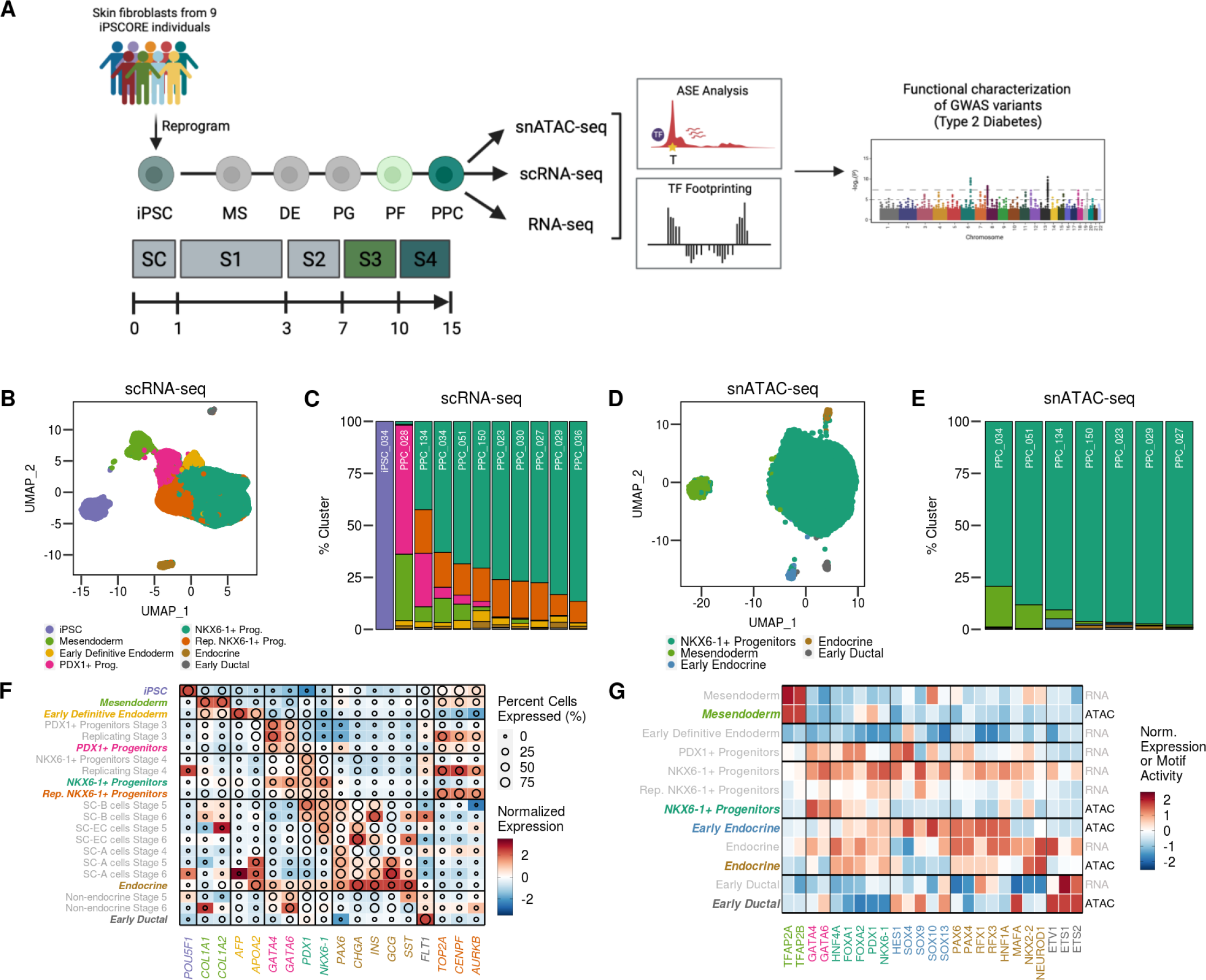
iPSC-PPC are largely comprised of NKX6-1+ progenitors. (A) Cartoon showing the overview design of the study. We differentiated iPSC-PPCs over a 15-day period and performed scRNA-seq, snATAC-seq, and bulk RNA-seq on matched samples and characterized regulatory variants for allele-specific effects on chromatin accessibility, transcription factor binding and gene expression. Using these profiles, we discovered variants that are active within iPSC-PPC regulatory elements and are associated with Type 2 Diabetes. (B) UMAP plot of scRNA-seq data from 83,871 single cells from one iPSC and ten iPSC-PPC samples. Each point represents a single cell color-coded by its assigned cluster. (C) Stacked bar plot showing the fraction of cells from each sample assigned to each cluster in scRNA-seq. Color-coding corresponds to the clusters in panel B. Differentiations PPC029 and PPC036 were from the same iPSC clone. (D) UMAP plot of snATAC-seq data from 25,654 single nuclei from seven iPSC-PPC samples. Each point represents a single nuclei color-coded by its assigned cluster. (E) Stacked bar plot showing the fraction of cells from each sample assigned to each cluster in snATAC-seq. Color-coding corresponds to the clusters in panel D. (F) Heatmap comparing the z-normalized expression of known marker genes between iPSC-PPC and cells from the reference ESC-PPC study. Color intensity corresponds to the mean z-normalized expression across all cell types, and the diameter corresponds to the fraction of cells expressing the markers above the threshold of 1% of the maximum expression value. Clusters labeled in italicized color correspond to the clusters in panel B. Clusters labeled in grey correspond to clusters identified in ESC-PPC scRNA-seq. (G) Heatmap comparing the z-normalized motif activity scores from chromVAR for pancreatic-associated transcription factors in the snATAC-seq clusters from panel D. Also shown are the normalized expression of the pancreatic-associated transcription factors in the scRNA-seq clusters from panel B. Clusters labeled in italicized color correspond to the snATAC-seq clusters in panel D. Clusters labeled in grey correspond to the scRNA-seq clusters in panel B.

### iPSC-PPC cell types characterized by scRNA-seq and snATAC-seq

To better understand the cellular heterogeneity of the iPSC-PPCs, we performed scRNA-seq on one iPSC clone and all ten iPSC-PPC samples (83,971 cells, 18,217 expressed genes), as well as snATAC-seq on seven of these samples (26,564 nuclei, 288,813 peaks). We integrated the scRNA-seq samples using Seurat (Butler et al., 2018) and detected eight distinct cell clusters (Figure 1B,C, Figure S2, Figure S3, Table S2). We found that the majority of the cells belonged to one cluster (52,014 cells, 61.94%). This observation was also reflected in snATAC-seq where a large proportion of nuclei (24,560 nuclei, 92.46%) belonged to a single cluster despite the fewer number of clusters (five) detected in snATAC-seq (Figure 1D,E, Figure S4, Table S4).

To annotate each cell type in scRNA-seq, we compared the expression levels of marker genes in each of the eight clusters with the expression levels in eight pancreatic progenitor cell types over four different stages during embryonic stem cell differentiation in the ESC-PPC reference study (Figure 1F, Table S3) (Veres et al., 2019). The Veres et al. study (Veres et al., 2019) included cell types corresponding to populations in the iPSC-PPCs (PDX1+ progenitors, NKX6-1+ progenitors, endocrine cells, and non-endocrine cells) as well as advanced cells (α and β cells) not represented in the iPSC-PPCs. For the cell types present in both studies, we identified clusters within the iPSC-PPCs that exhibit similar expression profiles as the corresponding cells in the ESC-PPCs: PDX1+ progenitors, which expressed the transcription factors *GATA4*, *GATA6*, and *PDX1,* but not *NKX6-1*, indicating that these cells are not yet fully committed towards pancreatic and beta cell differentiation (Xuan and Sussel, 2016); pancreatic progenitor cells (hereafter referred to as NKX6-1+ progenitors), which expressed both *PDX1* and *NKX6-1* and corresponded to the predominant cluster in scRNA-seq; endocrine cells which expressed both endocrine markers *PAX6* and *CHGA* and the pancreatic hormones *INS, GCG,* and *SST*; and non-endocrine cells, which we identified as precursors for ductal cells (referred to as early ductal), expressed endothelial marker *FLT1* (Pictet et al., 1972; Reichert and Rustgi, 2011). Similar to Veres et al., we identified a subcluster within the NKX6-1+ progenitors that expressed cell division markers (*TOP2A*, *CENPF*, *AURKB*), indicating that the cells in this cluster were replicating NKX6-1+ progenitors. We also identified primitive cells in the iPSC-PPCs not represented in the Veres et al. study including mesendoderm and early definitive endoderm, which expressed markers for early embryonic development (*COL1A1, COL1A2, AFP, and APOA2*) (Nowotschin et al., 2019; Saykali et al., 2019; Teo et al., 2015) and iPSCs, which expressed the stem cell marker *POUF51* and corresponded to the iPSC scRNA-seq sample. These results are consistent with the differentiation protocol used in Veres et al. (Veres et al., 2019), generating similar albeit more advanced cell types than the differentiation protocol we used to generate the iPSC-PPCs.

While the nine iPSC-PPC samples all consisted of multiple cell types, the vast majority of the scRNA-seq cells were PDX1+ progenitors, NKX6-1+ progenitors or replicating NKX6-1+ progenitors. To determine if our results reflected the differentiation efficiency measured by flow cytometry, we compared the fraction of NKX6-1+ progenitors and replicating NKX6-1+ progenitors, with the fraction of cells that stained double-positive for *PDX1* and *NKX6-1*. We found that these two independent measurements were highly correlated (R = 0.843, p = 0.00218; Pearson’s correlation, Figure S7A), indicating that scRNA-seq captured the variable differentiation efficiency observed in FACS.

To annotate the snATAC-seq clusters, we compared motif activity scores of pancreatic-associated transcription factors from chromVAR (Schep et al., 2017) with their gene expression levels from scRNA-seq (Figure 1G, Table S4). While most cell types could be identified in both scRNA-seq and snATAC-seq, we observed several differences (Figure 1, Figure S5, Figure S6, Table S5, Table S6). Both technologies identified mesendoderm (strong motif activity scores for TFAP2A, TFAP2B, (Raap et al., 2021; Wang et al., 2011)), NKX6-1+ progenitors (GATA4/6, HNF4A, FOXA1/2 PDX1, NKX6-1), endocrine (HNF1A, MAFA, NEUROD1, NKX2-2), and early ductal (ETV1, ETS1, ETS2) cell type populations. However, with scRNA-seq, we were able to distinguish clusters of early definitive endoderm cells and *PDX1+* progenitors, which could not be distinguished in snATAC- seq from *NKX6-1+* progenitors, which were the predominant cell type. Furthermore, snATAC-seq could not discriminate between replicating and non-replicating iPSC-PPC. However, when we compared the cell type fractions of NKX6-1+ progenitors in snATAC-seq with the fraction of early definitive endoderm, PDX1+, NKX6- 1+ progenitors, and replicating cells in scRNA-seq, the fractions were significantly correlated (R = 0.956, p = 0.000783, Figure S7B). Interestingly, we identified two distinct endocrine cell clusters using snATAC-seq, which differed in motif activity levels of early endocrine (SOX4, SOX10 and SOX13 (Lioubinski et al., 2003; Xu et al., 2015)) and late endocrine transcription factors (NKX2-2 and NEUROD1 (Doyle and Sussel, 2007; Itkin-Ansari et al., 2005; Mastracci et al., 2013)). Because the early endocrine cells also showed high motif activity levels for PAX and RFX (Figure 1G), which regulate endocrine differentiation, we determined that these cells are committed to endocrine lineage but have not yet fully developed into mature endocrine cells. Overall, while our results show that while scRNA-seq and snATAC-seq capture slightly different iPSC-PPC cell types, the predominant cell type across all samples in both assays consisted of NKX6-1+ progenitors.

### Endocrine cells express combinations of three pancreatic endocrine hormones

ESC-PPCs have been shown to produce polyhormonal endocrine cells (Shahjalal et al., 2018). We tested whether the 952 cells in the endocrine cluster expressed combinations of *INS* (insulin), *GCG* (glucagon), and *SST* (somatostatin, Figure S8). We observed that 50.7% of endocrine cells expressed *INS*, 18.7% expressed *GCG*, and 30.9% expressed *SST*. Of these cells, 23.7% expressed only one of the three hormones and 32.8% expressed at least two hormones, with *INS* and *SST* being the most common combination (16%) and 11% expressed all three hormones. While these hormones were also expressed in non-endocrine cell clusters, they were expressed in less than 5% of the cells. These results suggest that, while the protocol results in asynchronous differentiation, NKX6- 1+ progenitors, which are the most common cell type in the iPSC-PPCs, represent early pancreatic lineage to endocrine cells.

### Characterizing regulatory genetic variation in snATAC-seq

To understand the potential of iPSC-PPC as a model system to study pancreas regulatory genetics, we investigated the potential effects of variants overlapping 203,895 autosomal snATAC-seq peaks using three methods: 1) by identifying active transcription footprints and detecting their overlapping variants, using TOBIAS (Bentsen et al., 2020); 2) by predicting the allelic effects of variants in snATAC-seq peaks, using deltaSVM (Ghandi et al., 2014; Ghandi et al., 2016; Yan et al., 2021); and 3) by measuring allelic-specific effects (ASE) on heterozygous variants in snATAC-seq peaks.

To annotate accessible chromatin regions in iPSC-PPC, we identified 3,871,477 unique footprints for 746 transcription factors at 57,797 snATAC-seq peaks (28.3%) using TOBIAS (Bentsen et al., 2020; Stormo, 2013). We observed multiple footprints at the same peak for two reasons: 1) multiple transcription factors may bind to the same peak; and 2) since TOBIAS identifies transcription factor footprints independently for each transcription factor and determines the presence of a bound footprint based on each transcription factor motif, it cannot distinguish between transcription factors with highly similar motifs (for example: NKX6-1 and NKX6-2); therefore, multiple transcription factors in the same family may be identified as bound to the same footprint. To identify variants with potential effects on transcription factor binding in iPSC-PPC, we selected 325,942 common SNPs (≥ 5% minor allele frequency across 273 iPSCORE individuals (Panopoulos et al., 2017)) in 134,065 snATAC-seq peaks (65.8% of all peaks) and found that 35,248 (10.8%) overlapped at least one of the 3,871,477 active transcription factor footprints, corresponding to 19,775 peaks (9.7%) and 107,354 footprints (Table S7). Although TOBIAS identifies transcription factor footprints, indicating the genomic loci where a transcription factor is bound, it does not provide any information or prediction about the potential effects of the genotype of variants on transcription factor binding.

Next, to examine the allelic effects of variants in the snATAC-seq peaks we used recently published HT-SELEX data (Yan et al., 2021) generated for 94 transcription factors on ∼100,000 SNPs at T2D loci to train the deltaSVM model (Ghandi et al., 2014; Ghandi et al., 2016; Yan et al., 2021). This allowed us to use deltaSVM to predict the allelic effects of the 325,942 common SNPs on the binding of the 94 transcription factors in the 134,065 snATAC-seq peaks. We found that 52,653 SNPs (16.2%) in 42,511 peaks (20.8%) were predicted to overlap transcription factor binding sites and to have allelic effects on their binding (Table S7, Table S8). To validate these predictions, we investigated their overlap with the transcription factor footprints determined using TOBIAS for 89 transcription factors tested with both methods. For each of these transcription factors, we confirmed that variants predicted by deltaSVM to overlap bound transcription factor binding sites were more likely than expected to overlap the transcription factor footprint identified by TOBIAS (p = 2.2 x 10^-47^, t-test, Figure S9A, Table S9) and found a negative association between deltaSVM score and distance from each transcription factor footprint (p = 4.3 x 10^-15^, t-test, Figure S9B).

Finally, to measure allelic-specific effects in 48,738 snATAC-seq peaks that overlapped at least one SNP heterozygous in one or more of the seven tested samples (110,290 SNPs in total, including 86,660 of the ones tested with TOBIAS and deltaSVM, Table S7, Table S10), we utilized genotypes of the nine iPSCORE individuals from whole genome sequencing (DeBoever et al., 2017; Panopoulos et al., 2017). We divided SNPs according to whether they were heterozygous in one sample (termed “singletons” hereafter: 60,742 SNPs) or two or more samples (“multiplets”: 49,548 SNPs, Figure S10). We found 3,862 singleton and 3,487 multiplet SNPs with ASE (7,349 in total, FDR < 0.05, binomial test, adjusted using Benjamini-Hochberg’s method) in 5,583 peaks (2.25% of all peaks). We compared the allelic effects measured by ASE with the estimations by deltaSVM and observed a significantly positive correlation (R = 0.26, p-value = 3.0 x 10^-34^, Figure 2A). We also computed the correlation between snATAC-seq ASE and the deltaSVM predictions of allelic effects in each of these transcription factors. We observed a significantly positive correlation for 27 transcription factors (FDR ≤ 0.05, Benjamini-Hochberg’s method, Figure 2B, Table S11) and the distribution of correlation values across all transcription factors was significantly greater than zero (p = 3.7 x 10^-6^, t-test). The transcription factors with the strongest correlation included genes with known functions in embryonic development and pancreas, such as *JDP2* (Huang et al., 2011), *NFE2* (Kojayan et al., 2019), *ATF3* (Fazio et al., 2017), *CUX1* (Ripka et al., 2010) and *FOXB1* (Ma et al., 2016). We also observed a negative association between ASE measured on heterozygous variants in snATAC-seq peaks and their distance from transcription factor footprints (p = 6.45 x 10^-84^, t-test, Figure S9C, Table S12), indicating that variants with allelic effects are more likely than expected to affect transcription factor binding and that the results from the methods we employed to analyze the associations between genetic variation and chromatin function are concordant. Using three methods (TOBIAS, deltaSVM and ASE), we were able to detect or predict allelic effects or effects on transcription factor binding for 86,261 variants (Figure 2C). These results indicate that the genotypes of a large proportion of variants at iPSC-PPC snATAC-seq peaks are likely associated with altered binding affinities for transcription factors and therefore may be associated with adult pancreatic complex traits and disorders.

**Figure 2:**
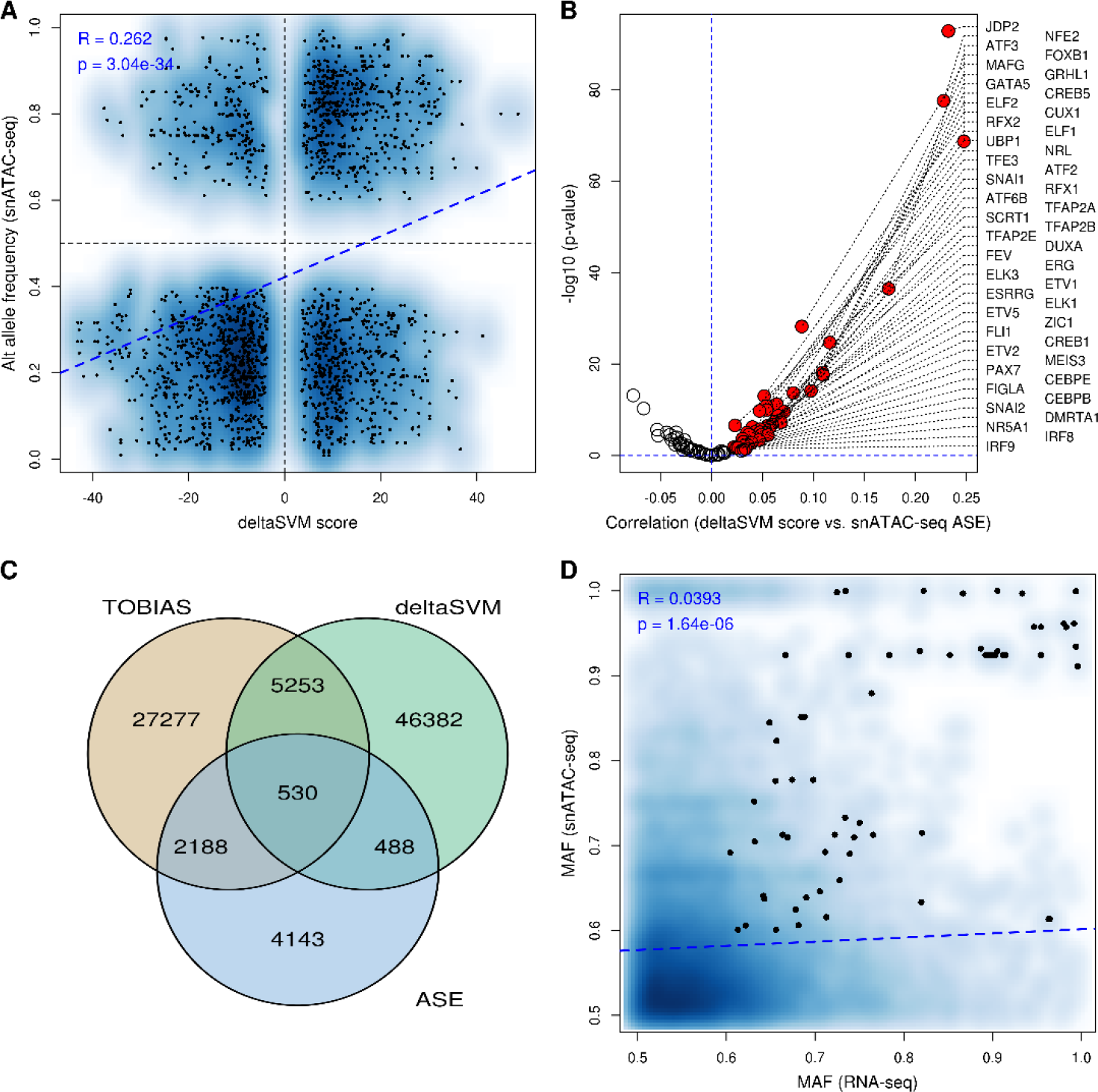
Allele-specific effects of SNPs in iPSC-PPC snATAC-seq peaks. (A) Scatterplot showing a significantly positive correlation (R = 0.26, p = 3.0 x 10^-34^) between SNPs predicted to have allelic effects across all 94 transcription factors measured by deltaSVM (X axis) and the alternative allele frequency (from snATAC-seq ASE analysis, Y axis). The blue dashed line represents the regression line. When considering all the SNPs tested for ASE and deltaSVM, the correlation was significantly positive (R = 0.019, p = 5.8 x 10^-80^), indicating that deltaSVM accurately predicts the allelic effects of SNPs in snATAC-seq on transcription factor binding. (B) Volcano plot showing the correlation between deltaSVM score and ASE measured by snATAC-seq. Each dot represents the correlation of all the SNPs for one of the 94 transcription factors tested by deltaSVM. Significant positively correlated transcription factors are highlighted in red and their names are indicated on the right. The volcano plot shows that the distribution of correlation values is significantly skewed towards positive values, confirming that, in general, deltaSVM predictions are concordant with snATAC-seq ASE values. (C) Venn diagram showing the overlap between the three methods (TOBIAS, deltaSVM and ASE) to characterize the evidence of functional effects (transcription factor binding or allelic effects) for each variant. (D) Smooth scatterplot showing a significant positive correlation (R = 0.039, p = 1.6 x 10^-6^) between the major allele frequency (MAF) of SNPs overlapping expressed genes, calculated by MBASED (X axis), and the MAF of SNPs overlapping snATAC-seq peaks at their corresponding promoters (Y axis). Dots represent genes with significant ASE that were associated with SNPs with ASE at their promoter. The blue dashed line represents the regression line.

To examine the correlation between ASE in iPSC-PPC regulatory elements due to genetic variation and to changes in gene expression, we performed bulk RNA-seq (scRNA-seq only has coverage at 3’ end of gene) for the seven samples with snATAC-seq and performed ASE on the transcriptome (Figure S10, Table S13). We observed a significant positive correlation between ASE at promoters and allelic bias with corresponding genes (R = 0.039, p = 1.6 x 10^-6^, Figure 2D). Although significant, the correlation between ASE at promoters and corresponding genes was weak, which is consistent with previous observations (Gate et al., 2018), and could reflect the fact that multiple proximal and distal regulatory elements can regulate the expression of a gene. Overall, these results show that regulatory genetic effects are consistent between the epigenome and the transcriptome.

### Regulatory variants with allelic effects in iPSC-PPC are associated with Type 2 Diabetes

To test if regulatory variants active in iPSC-PPC were associated with Type 2 Diabetes (T2D), we intersected the iPSC-PPC snATAC-seq peaks with the 380 99% functional credible sets (comprised of 66,607 SNPs) identified in a meta-analysis of 32 T2D GWAS from about 900,000 individuals (Mahajan et al., 2018). We found that the majority of credible sets (269, 70.8%) had at least one SNP (3,705 SNPs in total) that overlapped with iPSC-PPC snATAC-seq peaks (Table S14). These overlapping variants were more likely to have a higher rank (based on causality within a credible set, 8.6 x 10^-136^, Mann-Whitney U test) and a higher posterior probability of association (PPA) with T2D (p = 3.6 x 10^-68^) than SNPs outside of peaks (Figure 3A,B). These results indicate that regulatory elements active in iPSC-PPC are enriched for causal variants associated with T2D.

**Figure 3:**
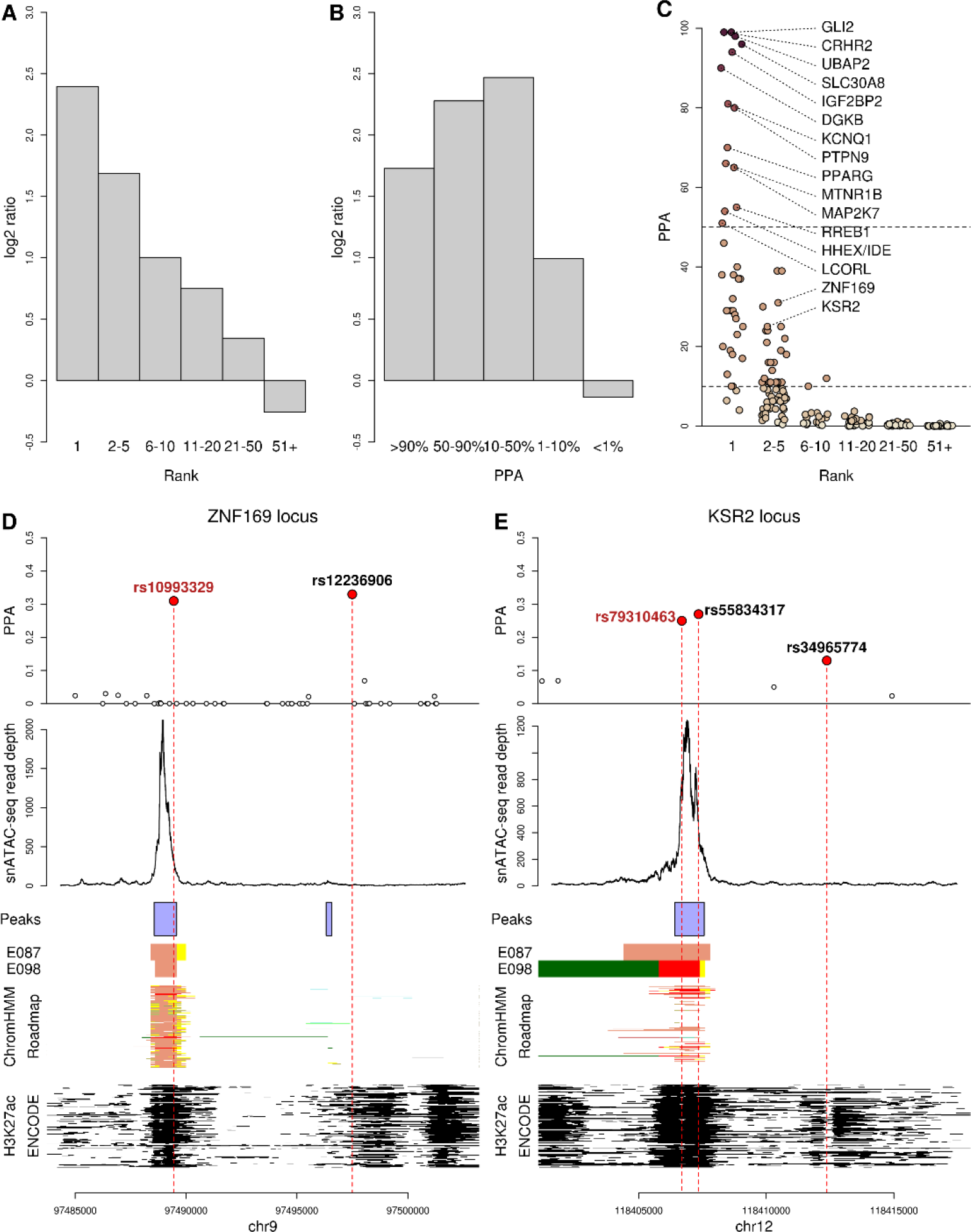
Associations between allele-specific effects and T2D-associated SNPs. (A, B) Barplots showing the enrichment for SNPs that overlap iPSC-PPC snATAC-seq peaks in T2D credible sets in different (A) rank or (B) PPA bins. X axis shows the rank or PPA bin and the Y axis represents the log2 ratio between the proportion of SNPs overlapping peaks and the proportion of SNPs not overlapping peaks. (C) Scatterplot showing the rank (X axis) and the PPA (Y axis: darker colors correspond to higher PPA values) for the 209 T2D loci with SNPs that display allelic effects or overlap transcription factor binding sites. The index genes associated with SNPs having PPA ≥ 50% and the index genes associated with the loci described in panels D and E are shown. Horizontal dashed lines represent PPA = 50% and PPA = 10%. (D, E) Two T2D loci (D: ZNF169; and E: KSR2) where the top-ranked SNP does not overlap snATAC-seq peaks or is not associated with ASE. The scatterplot (top) shows the PPA of each SNP included in the 99% credible set. The SNPs described in the text are shown and the SNPs with ASE are indicated in maroon. The second plot from the top shows the read depth from snATAC-seq. Below are shown: iPSC-PPC snATAC-seq peak coordinates (purple); 15 chromHMM chromatin states for two pancreas samples (E087: pancreatic islets; E098: pancreas) and 127 tissues included in Roadmap Epigenomics Program; and H3K27ac peak coordinates (in black) for 192 tissues obtained from the ENCODE data portal. Roadmap chromatin marks are colored as follows: 1) active TSS (red); 2) flanking active TSS (orange-red); 3) transcription at gene 5’ and 3’ (lime green); 4) strong transcription (green); 5) weak transcription (dark green); 6) genic enhancers (green-yellow); 7) enhancers (yellow); 8) ZNF genes and repeats (medium aquamarine); 9) heterochromatin (pale turquoise); 10) bivalent/poised enhancer or TSS (Indian red); 11) flanking bivalent/poised enhancer or TSS (dark salmon); 12) bivalent enhancer (dark khaki); 13) repressed polycomb (silver); 14) weak repressed polycomb (gainsboro); and 15) quiescent chromatin (white). In both examples, the SNPs with ASE overlap genomic regions that have H3K27ac peaks and are labeled as bivalent enhancers in many tissues. These examples show that the SNP with ASE, rather than the top-ranked SNP, is more likely to be functional and, therefore, causal for T2D.

To further characterize the potential of iPSC-PPC to study T2D genetics, we identified variants in the T2D credible sets that had high PPA or were the top-ranked and overlapped bound transcription factor binding sites (TOBIAS), had predicted allelic effects (deltaSVM) and/or validated allelic effects (ASE). Of the 269 credible sets with at least one SNP overlapping snATAC-seq peaks, 209 (77.7%) had at least one SNP potentially altering transcription factor binding affinities, including 163 with TOBIAS support, 152 with deltaSVM support and 65 with ASE. Of note, 73 had support from two methods and six from all three (Table S14). Among these 209 T2D loci, we found 65 SNPs that showed strong evidence of being causal, including 38 that were top-ranked and 27 that were not the top ranked but had high PPA (PPA > 10%, Figure 3C).

The 38 SNPs that were the top-ranked included six with PPA ≥ 90%, corresponding to the loci for *GLI2* (predicted to affect binding by deltaSVM and/or TOBIAS of HSF2, HSF4 and YY2), *CHCR2* (RFX1-5, SCRT1, SCRT2), *UBAP2* (HSF2 and HSF4), *SLC30A8* (PAX1 and PAX9), *IGF2BP2* (IRF1, PRDM1, ZNF384, and shows ASE in snATAC-seq) and *DGKB* (RFX1) and eight with PPA between 50% and 90%, corresponding to *KCNQ1* (ZNF148 and ZNF263, and shows ASE in snATAC-seq), *PTPN9* (HNF4A, HNF4G, NR4A1 and XBP1), *PPARG* (PPARG,RXRA and PROX1), *MTNR1B* (HSF2), *MAP2K7* (ZNF423), *RREB1* (ZBTB33), *HHEX/IDE* (ZNF460) and *LCORL* (ASCL1, ASCL2, BHLHE22, MYF5, MYOD1, MYOG, NHLH1, PTF1A, TCF12, ZBTB18 and ZSCAN29, Table S14). Many of the 209 potentially altered binding sites were associated with transcription factors with pancreatic functions: *HSF4* is expressed in pancreas and is associated with neuronal development (Nakai et al., 1997; Syafruddin et al., 2021); and YY2 is involved in the regulation of multiple cellular processes, including pluripotency and differentiation (Li et al., 2020); RFX3 is involved in pancreatic endocrine cells development (Ait-Lounis et al., 2007); SCRT1 is involved in the regulation of beta cell proliferation during differentiation(Sobel et al., 2021); IRF1 regulates the progression of pancreatic cancer (Sakai et al., 2014); ZNF148 is associated with pancreatic cancer risk (Fang et al., 2017); variants in HNF4A causes maturity-onset diabetes of the young and are associated with T2D (Yamagata, 2014); HNF4G is associated with glucose tolerance (Baraille et al., 2015); NR4A1 protects beta cells from apoptosis (Yu et al., 2015); XBP1 is required for the homeostasis of acinar cells (Hess et al., 2011); PPARG regulates multiple insulin-associated genes in beta cells (Gupta et al., 2010); RXRA negatively regulates glucose-stimulated insulin secretion (Miyazaki et al., 2010); PROX1 controls pancreas morphogenesis (Wang et al., 2005); PTF1A regulates acinar cell apoptosis (Sakikubo et al., 2018).

Twenty-seven credible sets had SNPs with allelic effects that were not the top ranked but had high PPA (PPA > 10%). These cases include the *ZNF169* locus, whose top-ranked SNP (rs12236906) has PPA = 33% but does not overlap any snATAC-seq peak, whereas its second-ranked SNP (rs10993329, PPA = 31%) overlaps a snATAC-seq peak and has deltaSVM predicted allelic effects (Figure 3D). Although rs12236906 has been indicated as the most likely causal SNP for this locus, our results suggest that rs10993329 is more likely to be functional. These observations are supported by the higher activity of the genomic region surrounding rs10993329 across multiple tissues (Figure 3D). We further investigated the predicted allelic effects of rs10993329 and found that it is associated with the loss of motifs for three members of the ETS family of transcription factors (ERG, FEV and FLI1), which play a role in pancreatic mesodermal development (Kobberup et al., 2007).

In the *KSR2* locus, the variants with the second- and third-highest PPA (rs79310463: 25%; and rs34965774: 13%) have both been described as causal for T2D and are both included in the GWAS Catalog (Buniello et al., 2019), whereas the variant with the highest PPA (rs55834317: 27%) is not associated with any GWAS. While rs79310463 and rs55834317 both overlap a snATAC-seq peak, only rs79310463 is associated with ASE and overlaps footprints for TFDP1 and ZNF263, suggesting that this SNP is more likely to have functional consequences in this locus (Figure 3E).

In other loci containing lower ranked SNPs (PPA > 10%), such as *HMG20A* and *IRS2*, multiple variants overlap iPSC-PPC snATAC-seq peaks and are predicted to have ASE, indicating that additional studies are needed to determine if multiple causal SNPs underlie the associations in these loci and whether they are functional. In conclusion, our genetic association analysis shows that many regulatory variants implicated in T2D are active and have allelic effects in iPSC-PPC, making these cells a suitable model system to identify and characterize the molecular mechanisms underlying T2D genetic associations.

## Discussion

In this study, we derived ten iPSC-PPC samples from nine unrelated individuals to generate matched scRNA-seq and snATAC-seq and determined that while the differentiation was asynchronous and similar to ESC-PPC (Veres et al., 2019), the derived cells largely consisted of a single cell type (NKX6-1+ progenitors). We characterized regulatory variants that overlapped open chromatin in iPSC-PPC and found that these variants are likely to have allelic effects on chromatin accessibility and may affect transcription factor binding. To validate the utility of iPSC-PPC to characterize and annotate genetic variants associated with adult T2D, we used previously fine-mapped 380 T2D risk loci (Mahajan et al., 2018). Enrichment analyses revealed that the majority of the T2D risk loci were located within open chromatin regions of iPSC-PPC. Furthermore, these loci contain variants that overlap active transcription factor binding sites and/or show allele specific effects on chromatin accessibility.

Our study identified 65 T2D risk loci containing SNPs with strong evidence of being causal (i.e. high PPA) that are associated with allelic-specific effects and/or predicted to affect transcription factor binding in the iPSC-PPCs. For 38 of the T2D risk loci, the top-ranked SNP (i.e. the SNP with highest PPA) had functional effects on transcription factor binding and/or chromatin accessibility; while in 27 loci, we observed that at least one lower-ranked SNP with high PPA (≥10%) was associated with allelic effects or altered transcription factor binding, suggesting that the top-ranked SNP is likely not causal. In certain cases, such as rs79310463 in the *KSR2* locus, the SNP we identified as being associated with allelic effects or transcription factor binding had been described as being causal in previous T2D studies (Ishigaki et al., 2020; Suzuki et al., 2019; Vujkovic et al., 2020); whereas in other loci (rs10993329 in the *ZNF169* locus), the SNP we predict to have functional effects had not previously been associated with T2D. Fine mapping using regulatory annotations, such as chromatin state maps in relevant tissues (Ernst and Kellis, 2012; Roadmap Epigenomics et al., 2015), prioritizes SNPs that overlap specific annotations (Pickrell, 2014); however, it is challenging to distinguish causal SNPs from variants that are in high LD with it. Here, we showed that characterizing the functional effects of individual SNPs using TOBIAS (Bentsen et al., 2020), deltaSVM predictions (Ghandi et al., 2014; Ghandi et al., 2016; Yan et al., 2021) or ASE, provides an alternative method to pinpoint the likely causal SNPs and may help discriminate neutral SNPs that are in high LD. Although the analyses proposed here identified SNPs that are associated with the active regulatory elements in iPSC-PPC, further analyses that integrate the results presented here with co-accessibility, expression quantitative trait loci (eQTLs) (Vinuela et al., 2020), chromatin accessibility QTLs (Alasoo et al., 2018), colocalization between QTLs and GWAS (Giambartolomei et al., 2014; Giambartolomei et al., 2018; Majumdar et al., 2018; Wallace, 2020) and, ultimately, experimental validation (Geusz et al., 2020), are needed to link their effects with their target genes and thus, functional mechanisms, as most regulatory elements are not in close proximity to promoters, and distal regulatory elements may regulate multiple genes (Oh et al., 2021). By empowering chromatin accessibility profiles with advanced tools such as transcription factor footprinting, allelic effect predictions, and co-accessibility, it is feasible to uncover novel molecular mechanisms that underlie the genetic risk of T2D.

Our study shows that by combining GWAS with epigenomic information from iPSC-PPC, it is feasible to gain insight into the molecular mechanisms underlying the associations between genetic variation and adult pancreatic complex traits and disease. Although we were able to determine the associations between SNPs in 209 T2D risk loci and transcription factor binding or allelic effects, the majority of the associations were predictions. Larger sample sizes would result in additional variants in ATAC-seq peaks and greater statistical power to test each variant-containing peak for ASE and downstream changes in gene expression. Indeed, to gain insight into global functional genetic variation, it will be necessary to obtain data for hundreds of iPSC-PPCs. With a small sample size, we show that iPSC-PPC provide a suitable model system to study the associations between genetic variation, regulatory mechanisms, and T2D, and that studies involving large numbers of samples could aid in the identification of causal variants at the majority of T2D risk loci.

## Methods

### iPSCORE subject information

We obtained 9 iPSC lines from the iPSCORE collection (Panopoulos et al., 2017) (Table S1). These lines were reprogrammed from skin fibroblasts collected from 9 unrelated subjects (8 female, 1 male), who ranged in age at time of donation from 21 to 65 years old, and represent three 1000 Genomes Project super populations: European American (7), Asian American (1), and African American (1). From each subject, whole blood samples were collected and used to generate and process whole genome sequence (WGS) data as previously described (D’Antonio et al., 2018; DeBoever et al., 2017). Briefly, reads were aligned against human genome b37 with decoy sequences (Genomes Project et al., 2015) using BWA-mem and default parameters (Li and Durbin, 2009). We applied the GATK best-practices pipeline for variant calling that includes indel-realignment, base- recalibration, genotyping using HaplotypeCaller, and finally joint genotyping using GenotypeGVCFs (DePristo et al., 2011; McKenna et al., 2010; Van der Auwera et al., 2013). The recruitment of these individuals was approved by the Institutional Review Boards of the University of California, San Diego and The Salk Institute (Project no. 110776ZF).

### iPSC-PPC Derivation

iPSC-PPCs were derived using STEMdiff™ Pancreatic Progenitor Kit (StemCell Technologies) following manufacture’s recommendations except as noted below. One iPSC line (from subject 90e8222f-2a97-4a3c-9517- fbd7626122fd) was independently differentiated twice (PPC_029 and PPC_036) resulting in a total of 10 derived iPSC-PPC samples.

*Expansion of iPSC:* One vial from each of 9 iPSC lines was thawed into mTeSR1 medium containing 10 μM ROCK Inhibitor (Selleckchem) and plated on one well of a 6-well plate coated with matrigel. iPSCs were grown until they reached 80% confluency and then passaged using 2mg/ml solution of Dispase II (ThermoFisher Scientific). To obtain a sufficient number of iPSCs for differentiation, iPSCs were passaged twice: 1) cells from the first passage were plated on three wells of a 6-well plate (ratio 1:3); and 2) cells from the second passage were plated on six wells of a 6-well plate (ratio 1:2).

*Monolayer plating (Day 0; D0)*: When the confluency of iPSC cells in the six wells of a 6-well plate reached 80%, cells were dissociated into single cells using Accutase (Innovative Cell Technologies Inc.). Single iPSC cells were resuspended at the concentration of 1.85 x10^6^ cells/ml in mTeSR containing 10μM ROCK inhibitor and plated on six wells of a 6-well. Cells were grown for approximately 16 to 20 hours to achieve a uniform 90- 95% confluency (3.7x10^6^ cells/well; about 3.9x10^5^ cells/cm^2^).

*Differentiation:* Differentiation of the confluent iPSC monolayers were initiated by the addition of STEMDiff Stage Endoderm Basal medium supplemented with Supplement MR and Supplement CJ (2ml/well) (D1). All following media changes were performed every 24 hours following initiation of differentiation (2ml/well). On D2 and D3, the medium was changed to fresh STEMDiff Stage Endoderm Basal medium supplemented with Supplement CJ. On D4, the medium was changed to STEMDiff Pancreatic Stage 2-4 Basal medium supplemented with Supplement 2A and Supplement 2B. On D5 and D6, the medium was changed to STEMDiff Pancreatic Stage 2-4 Basal medium supplemented with Supplement 2B. On D7, D8 and D9, the medium was changed to STEMDiff Pancreatic Stage 2-4 Basal medium supplemented with Supplement 3. On D10, D11, D12, D13 and D14, the medium was changed to STEMDiff Pancreatic Stage 2-4 Basal medium supplemented with Supplement 4.

*Harvest:* On D15 cells were dissociated using Accutase, collected and counted, and either processed fresh (scRNA-seq) or cryopreserved (scRNA-seq and snATAC-seq).

### Flow Cytometry

Each of the 10 iPSC-PPC differentiations were analyzed for co-expression of two pancreatic precursor markers, PDX1 and NKX6-1, using flow cytometry. Specifically, at least 2x10^6^ iPSC-PPC cells were fixed and permeabilized using the Fixation/Permeabilization Solution Kit with BD GolgiStopTM (BD Biosciences) following manufacturer recommendations. After the last centrifugation, cells were resuspended in 1X BD Perm/WashTM Buffer at the concentration of 1 x 10^7^/ml. For each flow cytometry staining, 2.5 x 10^5^ cells were stained with PE Mouse anti-PDX1 Clone-658A5 (BD Biosciences; 1:10) and Alexa Fluor® 647 Mouse anti- NKX6.1 Clone R11-560 (BD Bioscience; 1:10) or with appropriate class control antibodies, PE Mouse anti-IgG1 κ R-PE Clone MOPC-21 (BD Biosciences) and Alexa Fluor® 647 Mouse anti IgG1 κ Isotype Clone MOPC-21 (BD Biosciences). Cells were stained for 75 minutes at room temperature, washed three times, resuspended in PBS containing 1% BSA and 1% Formaldehyde, and immediately processed through FACS Canto II flow cytometer. FACS results were analyzed using FlowJo software V 10.4. The fractions of PDX1 and NKX6-1- positive cells varied across the analyzed iPSC-PPC lines, where percentages of PDX1/NKX6-1 double-positive cells ranged from 14.6 – 91.7% (mean = 60.0%; median = 71.0%).

### Generation of scRNA-seq

*Library Generation*: One 1 iPSC line (from subject: iPSC_PPC034) and 10 iPSC-PPC samples were used for scRNA-seq generation (Table S1). Fresh cells (i.e., not frozen) from the iPSC line and from seven iPSC-PPC samples were captured individually at D15. Four of these same iPSC-PPC samples were also captured as cryopreserved cells (immediately after thawing) along with three iPSC-PPC samples that were captured only as cryopreserved cells. Cells from four cryopreserved iPSC-PPC samples were pooled (RNA_Pool_1), and cells from the other 3 iPSC-PPC samples were pooled (RNA_Pool_2) prior to capture (Table S1). All single cells were captured using the 10x Chromium controller (10x Genomics) according to the manufacturer’s specifications and manual (Manual CG000183, Rev A). Cells from each scRNA-seq sample (1 iPSC, 7 fresh iPSC-PPCs, RNA_Pool_1, and RNA_Pool_2) were loaded on an individual lane of a Chromium Single Cell Chip B. Libraries were generated using Chromium Single Cell 3’ Library Gel Bead Kit v3 (10x Genomics) following manufacturer’s manual with small modifications. Specifically, the purified cDNA was eluted in 24μl of Buffer EB, half of which was used for the subsequent step of the library construction. cDNA was amplified for 10 cycles and libraries were amplified for 8 cycles.

*Sequencing*: Libraries produced from fresh and cryopreserved cells were sequenced on a HiSeq 4000 using custom programs (fresh: 28-8-175 Pair End and cryopreserved: 28-8-98 Pair End). Specifically, 8 libraries generated from fresh samples (1 iPSC and 7 iPSC-PPC samples) were pooled together and loaded evenly on 8 lanes and sequenced to an average depth of 163 million reads. Two libraries from seven cryopreserved lines (RNA_Pool_1 and RNA_Pool_2) were each sequenced on an individual lane to an average depth of 265 million reads. In total, we captured 99,819 cells. Figure S2 shows highly similar cell type proportions are observed in fresh and cryopreserved iPSC-PPCs.

### Processing scRNA-seq data

*Raw data processing.* We retrieved FASTQ files for 10 scRNA-seq samples (one iPSC, seven fresh iPSC-PPCs, one RNA_Pool_1, and one RNA_Pool_2) and used CellRanger V6.0.1 (https://support.10xgenomics.com/) with default parameters and v34lift37 (Harrow et al., 2012) gene annotations to generate single-cell gene counts and

BAM files for each individual sample (Table S1).

*Demultiplexing.* To reassign pooled cells iPSC-PPCs back to the original subject (RNA_Pool_1 and RNA_Pool_2; Table S1), we obtained the BAM files for each scRNA-seq sample and a VCF file containing SNPs (called from WGS) that are bi-allelic and located at UTR or exon regions on autosomes as annotated by Gencode v34lift37 (Harrow et al., 2012) calls from each of the nine subjects, two of which was not included in scRNA-seq pools but served as negative controls. The two files (BAM and VCF) were used as input to Demuxlet (Kang et al., 2018), which outputted the subject identities of each single cell based on genotype. We found that less than 1% of the cells mapped to negative controls after filtering for low quality cells and thus, were removed from downstream analyses.

*Data Processing.* To merge the 10 scRNA-seq samples (1 iPSC, 7 fresh iPSC-PPCs, RNA_Pool_1, and RNA_Pool_2), we first aggregated the samples that were sequenced as an independent batch (1 iPSC and 7 fresh iPSC-PPC) using the CellRanger V6.0.1 command *aggr* with no normalization. For each sample (aggregated sample, RNA_Pool_1, and RNA_Pool_2), we log-normalized the gene counts (*NormalizeData)* and identified the 2000 most variables genes using a threshold of 0.5 for the standardized log dispersion (*FindVariableFeatures*). We next applied Seurat’s standard integration workflow to adjust for batch differences between the samples. Specifically, we used *FindIntegrationAnchors* to identify a set of integration anchors between the samples using 30 dimensions computed from canonical correlation analysis (CCA). Next, we integrated the samples using *IntegrateData* and applied the standard downstream workflow of scaling the data (*ScaleData*), applying principle dimension reduction for 30 principle components (*RunPCA*), and then visualizing the single cells using Uniform Manifold Approximation and Projection (UMAP). To identify clusters, we used a shared-nearest-neighbor (SNN) graph of the significant PCs. To remove poor quality cells, we removed cells with fewer than 500 genes/cell or more than 50% of the reads mapping to the mitochondrial chromosome. We performed iterative clustering until all clusters driven by high mitochondrial reads or low number of genes were removed. Clusters with fewer than 250 cells were also removed. After filtering, 83,971 cells remained. We tested resolutions 0.05, 0.08, and 0.1 for clustering analyses and determined that 0.08 was more representative of the cell types predicted to be observed during stem cell differentiation into PPCs (Figure 1B,F, Figure S3).

### Annotation and validation of iPSC-PPC cells in scRNA-seq

To annotate the 83,971 iPSC-PPC cells, we used the expression of markers with known associations with pancreatic development and function, including *COL1A1*, *COL1A2* (mesendoderm) *AFP, APOA* (early definitive endoderm), *GATA4*, *GATA6, PDX1* (PDX1+ progenitors), *PDX1*, *NKX6-1* (NKX6-1+ progenitors), *PAX6, CHGA*, *INS*, *GCG*, *SST* (endocrine), *FLT1* (early ductal). We used *POU5F1* to identify the iPSC cluster. To obtain z-normalized expression values, we used cells with normalized expression values above 1% of the maximal expression, computed the average for each cluster, and then z-normalized across the 8 clusters. To validate cell type assignments, we used a reference scRNA-seq dataset from the ESC-B time course that captured cells from four differentiation stages (Veres et al., 2019): Stage 3 (Day 6; 7,982 cells), Stage 4 (Day 13; 6,960 cells), Stage 5 (Day 18; 4,193 cells), and Stage 6 (Day 25; 5,186 cells). Processed single cell gene counts and their associated metadata were downloaded from GEO (GSE114412). Z-normalized expression values were computed using the same procedure. We examined the transcriptome profiles at resolutions 0.05, 0.08, and 0.1, and found that the subclusters within the predominant cluster exhibited similar profiles to each other (Figure 1F, Figure S3, Table S2), confirming that they were NKX6-1+ progenitors. To identify differentially expressed genes, we performed Wilcoxon rank sum test between the normalized expression values of cells within the cluster and cells outside of the cluster (Table S3). P-values were adjusted using a Bonferroni correction, and genes with FDR ≤ 0.05 were considered differentially expressed.

### Generation of snATAC-seq

*Library Generation*: A total of 7 iPSC-PPC samples were used for snATAC-seq generation (Table S1). Cells from seven cryopreserved iPSC-PPCs samples were captured for snATAC-seq immediately after thawing. All seven samples have matched scRNA-seq. Cells from four cryopreserved iPSC-PPC samples were pooled (ATAC_Pool_1) and cells from the other 3 iPSC-PPC samples were pooled (ATAC_Pool_2) prior to capture (Table S1). Nuclei from two pools were isolated according to the manufacturer’s recommendations (Manual CG000169, Rev B), transposed, and captured as independent samples according to the manufacturer’s recommendations (Manual CG000168, Rev B). All single nuclei were captured using the 10x Chromium controller (10x Genomics) according to the manufacturer’s specifications and manual (Manual CG000168, Rev B). Cells for each sample were loaded on the individual lane of a Chromium Chip E. Libraries were generated using Chromium Single Cell ATAC Library Gel & Bead Kit (10x Genomics) following manufacturer’s manual (Manual CG000168, Rev B). Sample Index PCR material was amplified for 11 cycles.

*Sequencing*: Libraries were sequenced using a custom program (50-8-16-50 Pair End) on HiSeq 4000. Specifically, two libraries from seven cryopreserved iPSC-PPC samples (ATAC_Pool_1 and ATAC_Pool_2) were each sequenced on an individual lane.

### Processing snATAC-seq data

*Raw data processing.* For two snATAC-seq samples (ATAC_Pool_1 and ATAC_Pool_2; Table S1), we retrieved FASTQ files and used CellRanger V2.0.0 (https://support.10xgenomics.com/) to align files to the hg19 genome using *cellranger-atac count* with default parameters. NarrowPeaks were called using the MACS2 command *macs2 callpeak --keep-dup all --nomodel --call-summits* (Feng et al., 2012) on the BAM files merged from the two pooled samples and detected 288,813 peaks. Peaks called on ambiguous chromosomes or the mitochondrial genome were removed, leaving 280,079 peaks remaining. Using these peaks, each snATAC-seq sample was reanalyzed using *cellranger-atac reanalyze* to generate single-nuclei peak counts for each sample. To integrate the two snATAC-seq datasets for downstream analyses, we performed Signac integration (Butler et al., 2018) by first applying normalization (*RunTFIDF*) and linear dimensional reduction (*FindTopFeatures* and *RunSVD*) on each sample dataset. We then identified a random subset of 20,000 peaks and computed a set of integration anchors between the samples (*FindIntegrationAnchors* for 2,000 anchors) The two snaATAC-seq was integrated using *IntegrateData* and 2-30 most significant dimensions calculated from dimension reduction analyses. Finally, on the integrated dataset, dimension reduction was applied (*RunSVD* for 30 singular values), and single cells were visualized using UMAP (*RunUMAP* on 2:30 dimensions). Clusters were identified using a SNN-graph method using *FindNeighbors* and *FindClusters*. To remove low quality cells, we removed cells that satisfy one of the following criteria: 1) the number of peak region fragments < 2,000 or > 20,000, 2) the percentage of reads in peaks < 40%, 3) nucleosome signal > 1.5, or 4) TSS enrichment score < 2.5. Furthermore, we removed cells that do not visually belong to a cluster (i.e. cells that are scattered between two distinct clusters). We performed iterative clustering until we do not observe significant outliers of single cells. After filtering, 25,564 nuclei remained and clustering resolutions of 0.1, 0.15, and 0.2 were tested.

*Demultiplexing.* To reassign pooled nuclei back to the original subject from two snATAC-seq samples

(ATAC_Pool_1 and ATAC_Pool_2; Table S1), we applied Demuxlet (Kang et al., 2018) to the two samples using the same set of reference variants as stated above.

### Annotation of iPSC-PPC nuclei in snATAC-seq using chromVAR

To determine the cell types within the integrated snATAC-seq dataset, we used chromVAR (Schep et al., 2017) within the Signac pipeline to identify transcription factor motifs from the JASPAR 2020 database (Fornes et al., 2020)that are enriched for accessible chromatin for each cluster. Specifically, we used the *RunChromVAR* function in Signac and the hg19 reference (BSgenome.Hsapiens.UCSC.hg19) to compute a deviation z-score for each motif in each cell. To annotate the cell types, we examined the motif activities of transcription factors with known developmental or pancreatic functions: TFAP2A/B (mesendoderm), GATA4/6 (PDX1+ progenitors), HNF4A, FOXA1/2, PDX1, NKX6-1 (NKX6-1+ progenitors), HES1, SOX4/9/10/13 (early endocrine), PAX4/6, RFX1/3, HNF1A, MAFA, NKX2-2, NEUROD1 (endocrine), and ETV1, ETS1, ETS2 (early ductal). To validate our annotations, we compared the motif activities to their gene expression in scRNA-seq using the same z-normalization method. We examined the motif activity profiles at resolutions 0.1, 0.15, and 0.20 (Figure 1G,

Figure S4, Table S4), and reasoned that because subclusters within the predominant cluster expressed both PDX1 and NKX6-1 but at varying levels, we collapsed these clusters into NKX6-1+ progenitors. Resolution 0.1 was used for downstream analyses. To identify differentially expressed peaks, we applied the *FindAllMarkers* function in Signac with default parameters. Peaks with FDR ≤ 0.05 were considered differentially expressed (Table S5).

### chromVAR motif enrichment analyses

To identify differentially enriched motifs for each snATAC-seq cluster at resolutions of 0.1, 0.15, and 0.20, we computed a Wilcoxon rank sum test for each motif and cell type cluster, comparing the chromVAR z-score distributions of a random sample of 2,000 cells within the cluster and a random sample of 2,000 cells outside of the cluster. Then, for each cluster, we applied a Bonferroni correction to account for multiple testing. Motifs with FDR ≤ 0.05 were considered differentially enriched (Table S6).

### Processing transcription factor footprints using TOBIAS

To characterize regulatory variants for transcription factor binding sites, we used TOBIAS (Bentsen et al., 2020) to identify the binding sites of transcription factors. We merged BAM files from snATAC_Pool_1 and snATAC_Pool_2 and corrected for bias from Tn5 cutsites using TOBIAS function *ATACorrect*. Using the cutsite tracks from *ATACorrect*, we computed footprint scores across the regions using *FootprintScores*. Using the footprint scores along with transcription factor binding motifs from JASPAR2020 (Fornes et al., 2020), we then estimated the binding positions of each transcription factor footprint across the genome. Using these positions, we calculated the distance of each of the 325,942 SNPs in snATAC-seq peaks from its closest transcription factor footprint on the same peak using *bedtools closest -d*. Information about the footprints can be found in Table S7.

### Prediction of allelic effects using deltaSVM

We obtained deltaSVM (Ghandi et al., 2014; Ghandi et al., 2016) models on 94 transcription factors from the Genetic Variants Allelic TF Binding Database (GVATdb) (Yan et al., 2021). As training sets for the deltaSVM models, GVATdb includes the results from a SNP-SELEX experiment that analyzed the allelic effects of 95,886 noncoding variants located in close proximity with 110 T2D loci on transcription factor binding. Although GVATdb investigated 533 transcription factors, only 94 were associated with high-confidence deltaSVM models (Yan et al., 2021) and were used in this study. We selected 325,942 SNPs with at least 5% minor allele frequency in the iPSCORE cohort (Panopoulos et al., 2017) that overlapped the 203,895 snATAC-seq peaks using bcftools (Danecek et al., 2021). We found that 134,065 peaks had overlapping SNPs. We ran the deltaSVM pipeline developed within GVATdb (https://github.com/ren-lab/deltaSVM) on each of these variants. This resulted in 30,638,548 tests (325,942 SNPs by 94 transcription factors). To detect SNPs with a predicted allelic effect on transcription factor binding, we filtered these tests based on *seq_binding == “Y”* and *preferred_allele != “None”*. *seq_binding* refers to whether the transcription factor is predicted to be bound to the locus overlapping the tested SNP and preferred_allele describes whether the SNP is associated with improved binding affinity for the transcription factor (“*Gain*”), decreased binding affinity (“*Loss*”) or is not associated with changes in transcription factor binding affinity (“*None*”). We found 84,196 tests passing these filters, for a total of 52,653 unique SNPs, as one SNP may be predicted to affect the binding affinity for more than one transcription factor.

### Allele-specific effects in snATAC-seq peaks

We filtered the total 288,813 snATAC-seq peaks to include peaks on autosomal chromosomes, with MACS2 score ≥ 100, and outside of ENCODE blacklist regions (hg19-blacklist.v2.bed.gz) (Amemiya et al., 2019; Consortium, 2012). We intersected the resulting 203,895 peaks with the SNPs heterozygous in at least one of the seven samples that underwent snATAC-seq (Figure S10). Variant information was obtained from our previously published whole genome sequencing (dbGaP: phs001325) (DeBoever et al., 2017; Jakubosky et al., 2020a; Jakubosky et al., 2020b). We obtained 110,290 SNPs with read depth ≥ 20 in at least one heterozygous sample, which we divided into: 1) 60,742 SNPs heterozygous only in one sample (singletons); and 2) 49,548 SNPs heterozygous in multiple samples (multiplets). We calculated allele-specific effects (ASE) for each SNP independently in each heterozygous sample using a two-sided binomial test with the alternative hypothesis that both alleles were equally likely to be observed (p = 0.5). We further removed all multiplets with inconsistent effects: only 22,717 multiplet SNPs (61,562 tests) having the sign of 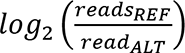 consistent across all heterozygous samples were retained for downstream analyses. FDR correction was performed on a per-sample basis using Benjamini-Hochberg’s method on all singleton and multiplet SNPs passing this filter (17,199 – 36,517 tests). To test for the correspondence between ASE in snATAC-seq and bulk RNA-seq, we obtained the coordinates of Genecode v34lift37 (Harrow et al., 2012) promoters and intersected them with coordinates of snATAC-seq peaks displaying ASE using *bedtools intersect*. We obtained 826 and 759 genes whose promoters overlapped singleton and multiplet SNPs, respectively. Of these genes, we retained 447 and 497, respectively, that were expressed (TPM > 1 in the heterozygous samples) and that had read depth ≥ 10 for at least one heterozygous variant.

### Generation of bulk RNA-seq

*Library generation and sequencing:* For 10 iPSC-PPC samples, RNA was isolated from total-cell lysates using the Quick-RNA^TM^ MiniPrep Kit (Zymo Research) with on-column DNAse treatments. RNA was eluted in 48ul RNAse-free water and analyzed on a TapeStation (Agilent) to determine sample integrity. All iPSC-PPC samples had RNA integrity number (RIN) values between 9.4 and 10. Illumina TruSeq Stranded mRNA libraries were prepared and sequenced on HiSeq 4000 to an average depth of 58.6 M 100-bp paired-end reads per sample.

*Raw data processing.* FASTQ files were obtained for the 10 iPSC-PPC samples and processed with a similar pipeline used in our previous studies (D’Antonio-Chronowska et al., 2019; D’Antonio et al., 2021; DeBoever et al., 2017). Briefly, RNA-seq reads were aligned with STAR (2.7.3) (Dobin et al., 2013) to the hg19 reference using Gencode v34lift37 (Harrow et al., 2012) splice junctions with default alignment parameters and the following adjustments: *-outFilterMultimapNmax 20, -outFilterMismatchNmax 999, -alignIntronMin 20, - alignIntronMax 1000000, -alignMatesGapMax 1000000*. Bam files were sorted by coordinates using Samtools (1.9.0), and duplicate reads were marked using Samtools (1.9.0) (Danecek et al., 2021). TPM values were estimated from STAR transcriptome bam file using RSEM (1.2.20) (Li and Dewey, 2011). RNA-seq QC metrics were collected from Samtools (1.9.0) flagstat and idxstats and/or Picard (2.20.1) *CollectRnaSeqMetrics* (2019).

*Sample quality control.* To confirm the subject identify assigned to each bulk RNA-seq, we tested common variants from the 1000 Genomes Phase 3 panel (Genomes Project et al., 2015)that are bi-allelic and have minor allele frequency between 45% to 55%. For each sample, genotype likelihoods were estimated using BCFtools (Danecek et al., 2021) (1.9.0) mpileup relative to the hg19 reference, and genotypes were called using BCFtools (1.9.0) *call*. Genotypes were filtered by a threshold of 10 for total read depth. Identity-by-state (IBS) was then estimated with PLINK (Purcell et al., 2007) *genome* for each pairwise comparison between the inferred genotypes from RNA-seq and the genotypes from WGS. RNA-seq samples were correctly matched with the subjects based on the highest pihat for each RNA-seq and individual pair.

### Allele-specific effects in bulk RNA-seq using MBASED

To detect the allele-specific effects of gene expression in bulk RNA-seq from seven samples (7 subjects), we used an R package MBASED (Mayba et al., 2014), which uses a meta-analysis approach that aggregates information from all SNPs within the gene body to measure gene-level ASE. For ASE analysis, we only considered genes on autosomes and that is expressed. We determined a gene to be expressed if the gene is expressed with at least 1 TPM in at least 10% of the seven samples. Of the 62,492 genes, 18,217 genes (29.2%) were expressed, of which 10,715 were on autosomes. For each sample and gene, we obtained read counts for the reference and alternate allele using *samtools* mpileup at the SNP loci for which the sample was heterozygous. We ran the 1-sample analysis on MBASED, obtained the major allele frequency and p-value of ASE for each gene, and applied multiple test correction using Benjamini-Hochberg’s method. As default, MBASED removed genes with less than 10 reads for read depth. We determine a gene to display ASE if FDR ≤ 0.05 and major allele frequency ≥ 0.6.

### Processing T2D loci

We obtained the genomic coordinates of each of 380 fine-mapped T2D loci from Mahajan et al. (Mahajan et al., 2018) . For each SNP with PPA > 0 in each locus, we extracted all its iPSC-PPC snATAC-seq peaks using *bedtools intersect* (Quinlan and Hall, 2010).

### Obtaining Roadmap and ENCODE epigenomic data

We downloaded 15 chromHMM chromatin state annotations (Ernst and Kellis, 2012) in 127 tissues included in the Roadmap Epigenomics Program (Roadmap Epigenomics et al., 2015). Each state was predicted using a hidden Markov model (HMM) on the signal from five histone modification ChIP-seq experiments, including H3K4me3, H3K4me1, H3K36me3, H3K9me3 and H3K27me3. We obtained BED file from https://egg2.wustl.edu/roadmap/web_portal/chr_state_learning.html.

We downloaded H3K27ac peak coordinates for 192 tissues from the ENCODE data portal (https://www.encodeproject.org/) (Consortium, 2012; Davis et al., 2018). We selected all H3K27ac samples with NarrowPeak BED files that passed all quality filters established by ENCODE.

## Data availability

iPSC-PPC scRNA-seq and snATAC-seq data was submitted to GEO: <u>GSE152610</u> (token *khyrckqqzpsprib*).

Seurat objects, including snATAC-seq and scRNA-seq, results from differential gene expression, differential peak expression and motif enrichment analysis have been deposited to Figshare: https://figshare.com/projects/Regulatory_variants_active_in_iPSC-derived_pancreatic_progenitor_cells_are_globally_associated_with_Type_2_Diabetes_in_adults/119706. ESC-PPC reference scRNA-seq was obtained from GEO (<u>GSE114412</u>). ChromHMM chromatin states were obtained from https://egg2.wustl.edu/roadmap/web_portal/chr_state_learning.html. H3K27ac peak coordinates were obtained <u>from</u> https://www.encodeproject.org/. Fine mapped T2D loci were obtained from the Diagram Consortium (https://www.diagram-consortium.org/).

## Acknowledgements

This work was supported in part by a California Institute for Regenerative Medicine (CIRM) grant GC1R-06673 and NIH grants HG008118, HL107442, DK105541, and DK112155. JPN, TDA and MKRD were supported by the National Library of Medicine Training Grant T15LM011271. BS was supported by CIRM Bridges Stem Cell Research and Therapy Training Grant, Award #EDUC2-08375. We would like to thank Drs. Maike Sander, Bing Ren and Kyle Gaulton for insightful scientific conversations.

## Author information

KAF, ADC, and MD conceived the study. ADC, KF and BMS performed iPSC-PPC differentiations. ADC generated the molecular data. JPN, MKRD, HM, MD, and TDA performed computational analysis. JPN and MKRD performed scRNA-seq data processing and analyses. HM, MD and JPN performed the snATAC-seq data processing and analyses. KAF oversaw the study. KAF, JPN, MKRD and MD prepared the manuscript.

## iPSCORE Consortium

University of California, San Diego, La Jolla, CA 92093, USA

Angelo D. Arias, Timothy D. Arthur, Paola Benaglio, Matteo D’Antonio, Agnieszka D’Antonio-Chronowska, Christopher DeBoever, Margaret K.R. Donovan, Kelly A. Frazer, Olivier Harismendy, David Jakubosky, Kristen Jepsen, He Li, Hiroko Matsui, Naoki Nariai, Daniel T. O’Connor, Jennifer P. Nguyen, Fengwen Rao, Erin N. Smith, William W. Young Greenwald

Salk Institute for Biological Studies, La Jolla, CA 92037, USA

Athanasia D. Panopoulos, W. Travis Berggren, Kenneth E. Diffenderfer

## SUPPLEMENTAL FIGURES

**Figure S1:**
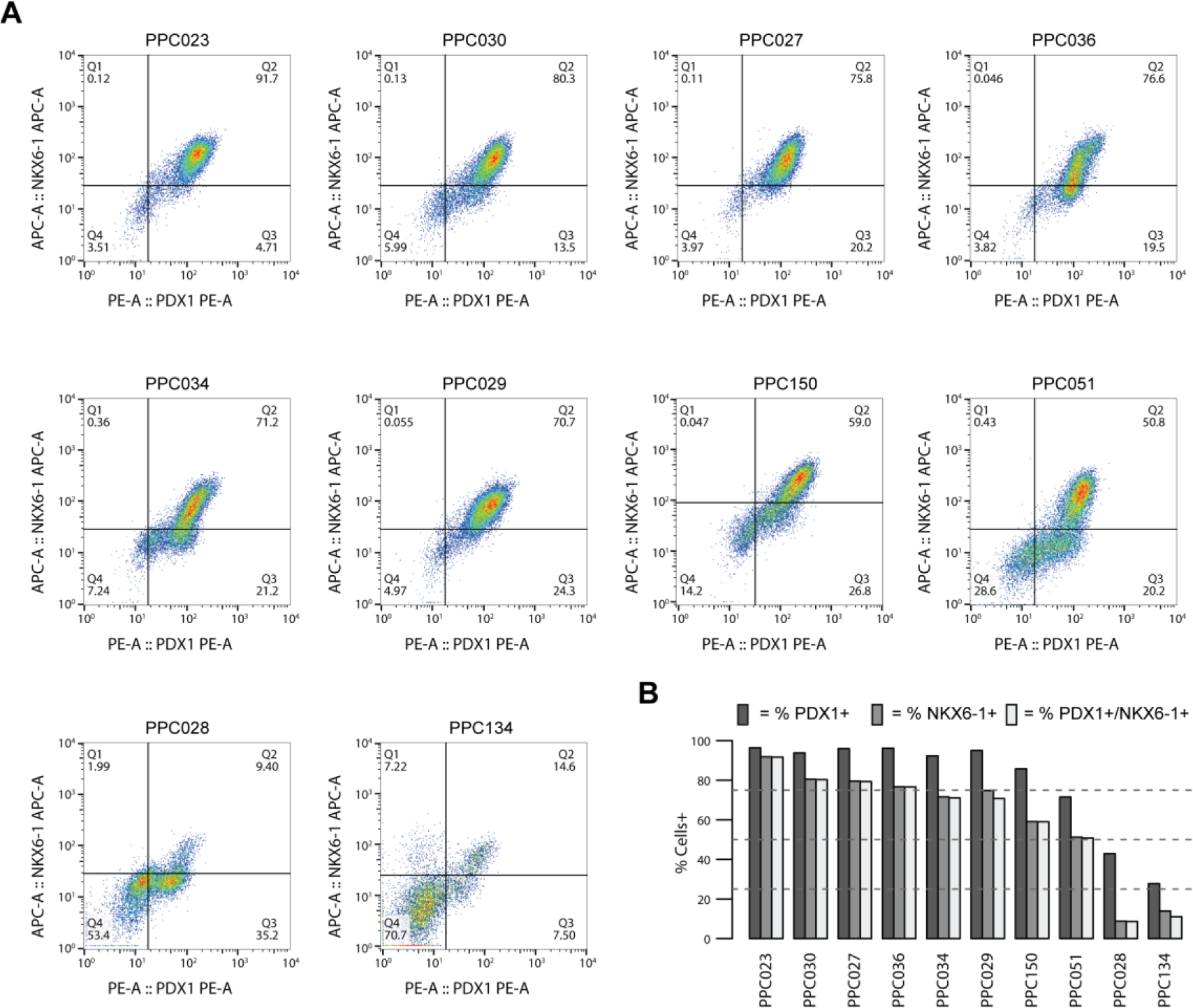
Measurement of PDX1-postive and NKX6-1-positive cells by flow cytometry. (A) Flow cytometry analysis at D15 of ten iPSC-PPC differentiations. The fraction of cells stained for PPC markers, PDX1 and NKX6-1, were measured. Differentiations PPC029 and PPC036 were from the same iPSC clone. (B) Bar plot showing the fraction of iPSC-PPC cells positively stained for PPC markers, *PDX1* and *NKX6-1*, and positive for both *PDX1* and *NKX6-1*. Differentiations PPC029 and PPC036 were from the same iPSC clone.

**Figure S2:**
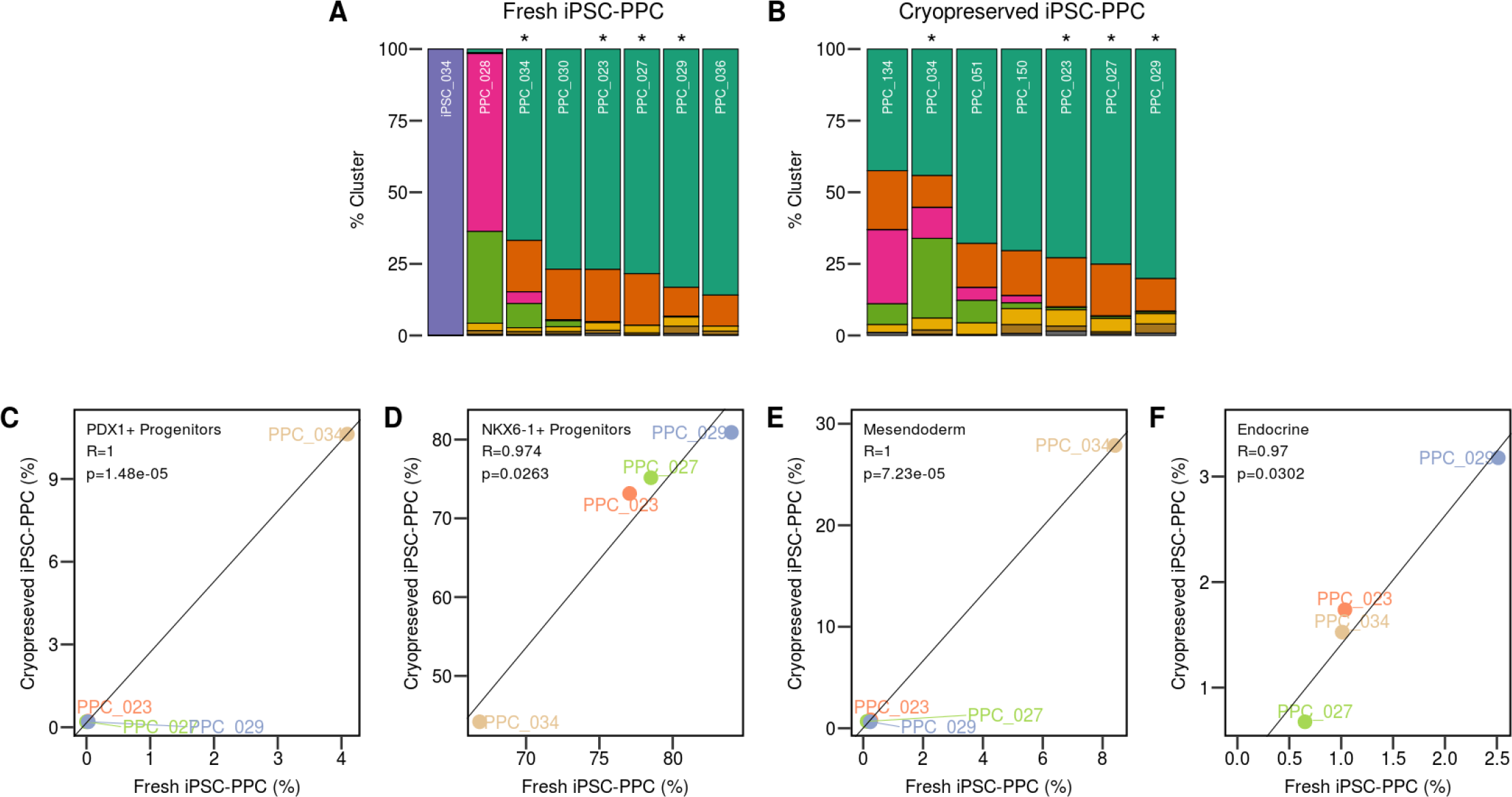
Similar cell type proportions between fresh and cryopreserved cells. To determine if cryopreservation influences our detection of iPSC-PPC cell types, we integrated scRNA-seq obtained from eight fresh (i.e., not frozen) sample preparations (seven iPSC-PPC and one iPSC sample, aggregated into a single pool) with two pools of four and three cryopreserved iPSC-PPC samples. To assign the sample identity of each cell in the two cryopreserved pools, we performed sample deconvolution with demuxlet using genotype information from whole genome sequencing of nine individuals (DeBoever et al., 2017), two of which served as negative controls (i.e. samples from these two individuals were not included in the cryopreserved pools, Table S1). For the fresh preparations, each sample was processed independently from the others, therefore deconvolution using Demuxlet (Kang et al., 2018) was not required. For each of the four iPSC-PPC samples with matched fresh and cryopreserved preparations (indicated by the asterisks above the bar plots), we compared the proportions of cells in PDX1+ progenitors, NKX6-1+ progenitors, mesendoderm, and endocrine, and found that fresh and cryopreserved samples were highly correlated. These results indicate that fresh and cryopreserved cells can be used interchangeably to characterize the cellular composition of iPSC-PPC. The figure shows: (A) Stacked bar plots showing the fraction of cells from each fresh iPSC-PPC sample assigned to each cell type using the same color coding as Figure 1B. Asterisks indicates the four iPSC-PPC samples with matched fresh and cryopreserved preparations. (B) Stacked bar plots showing the fraction of cells from each cryopreserved iPSC-PPC sample assigned to each cell type using the same color coding as Figure 1B. (C-F) For the four iPSC-PPC samples with matched fresh and cryopreserved samples, we show the association between each preparation by comparing the fraction of cells for (C) PDX1+ progenitors, (D) NKX6-1+ progenitors, (E) mesendoderm and (F) endocrine.

**Figure S3:**
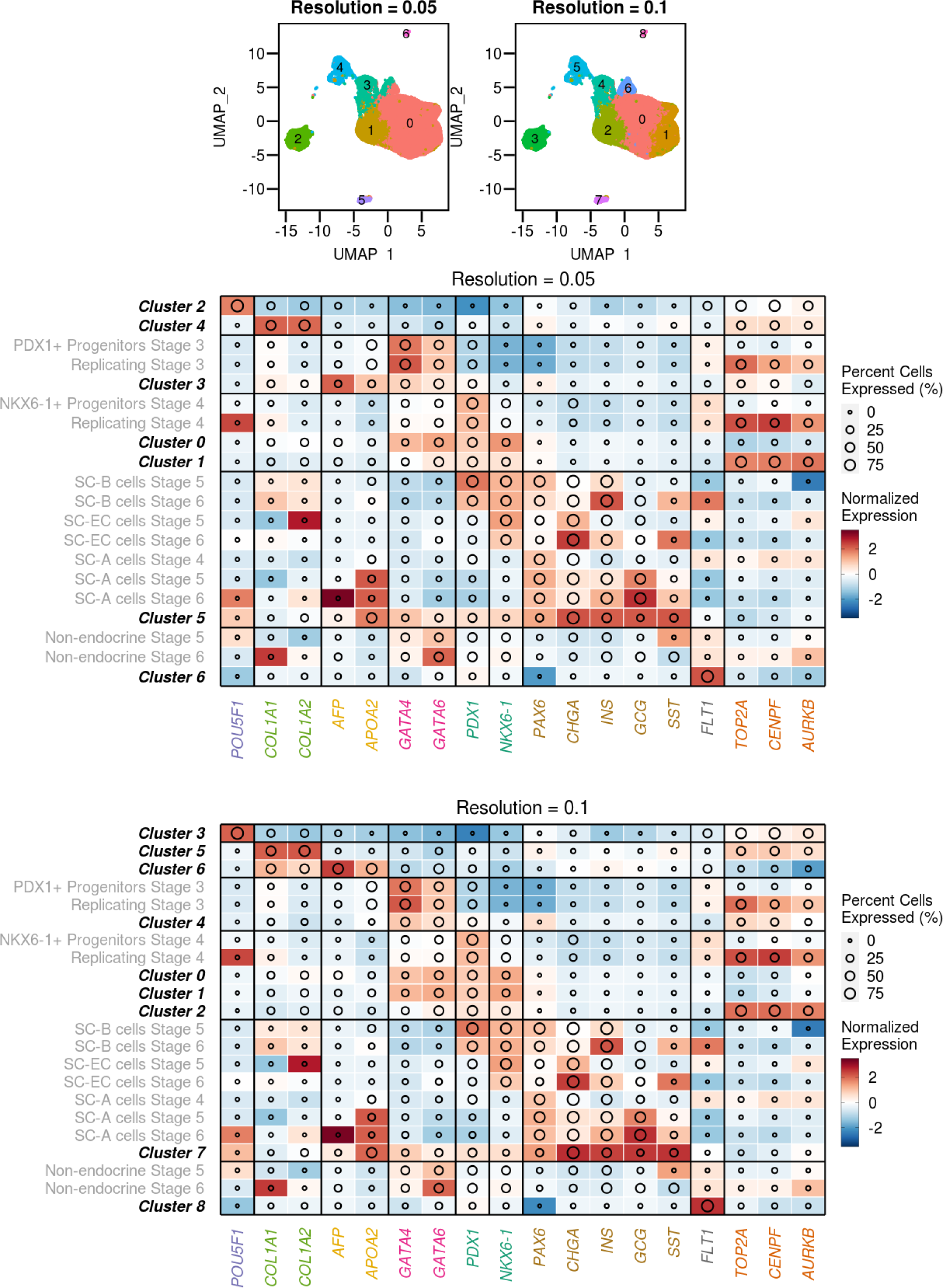
Clustering of iPSC-PPC scRNA-seq at two additional resolutions. Clustering was performed on 83,971 single cells from one iPSC and ten iPSC-PPC samples at resolutions 0.05, 0.08 (shown in Figure 1B) and 0.1. With increasing resolution, we found that cluster 0 (*NKX6-1+* progenitors) was further divided into subclusters. While all subclusters expressed *PDX1* and *NKX6-1*, one cluster expressed cell division markers (*TOP2A*, *CENPF*, and *AURKB*) at high levels, indicating that this cluster consists of replicating NKX6-1+ progenitors. Because the expression profiles are similar between the subclusters within cluster 0 at resolution 0.1, we used resolution 0.08 for downstream analyses where the subclusters were collapsed to form NKX6-1*+* progenitors (Figure 1B). Red-to-blue shade in the heatmaps indicates z-normalized expression and the diameter represents the fraction of cells that express 1% of the maximal expression within that cell type. Cluster labels in black correspond to clusters in iPSC-PPC scRNA-seq (shown in the UMAP plots above the heatmaps). Cluster labels in grey correspond to clusters identified ESC-PPC scRNA-seq (Veres et al., 2019).

**Figure S4:**
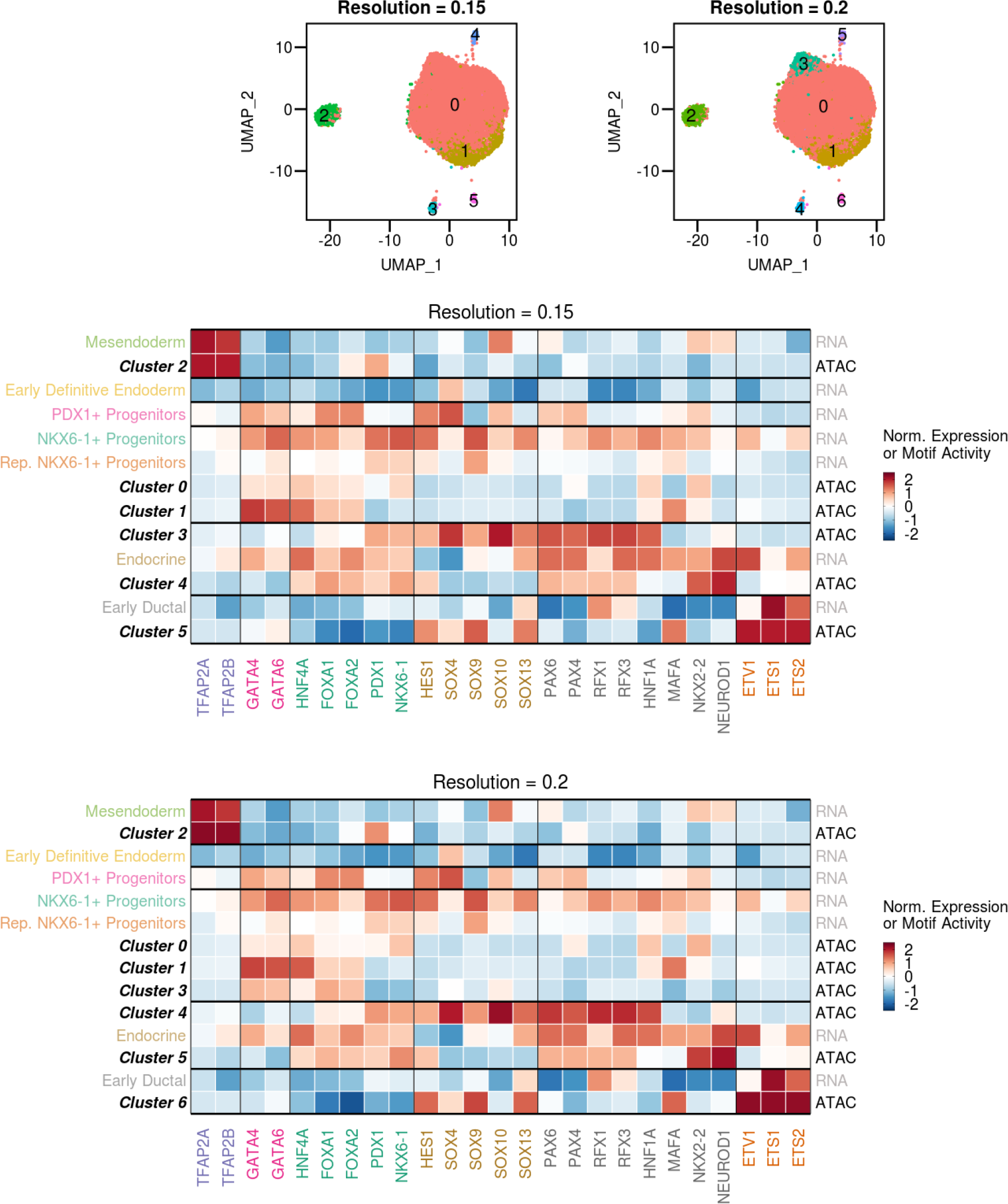
Clustering of iPSC-PPC snATAC-seq at two additional resolutions. We performed clustering analyses on 26,564 single nuclei from seven iPSC-PPC samples at resolutions 0.1 (shown in Figure 1D), 0.15 and 0.2. At resolutions 0.1, 0.15 and 0.20, we identified a total of five, six and seven clusters respectively. With increasing resolution, cluster 0 was further divided into subclusters that largely consist of *NKX6-1+* progenitors but with varying levels of PDX1 and NKX6-1 motif activities. Cluster labels in black correspond to clusters in snATAC-seq described in the UMAP plots above the heatmaps. Colored cluster labels correspond to iPSC-PPC scRNA-seq clusters described in Figure 1B. Red-to-blue shade in the heatmaps indicates Z-normalized expression for scRNA-seq or chromVAR motif activity for snATAC-seq.

**Figure S5:**
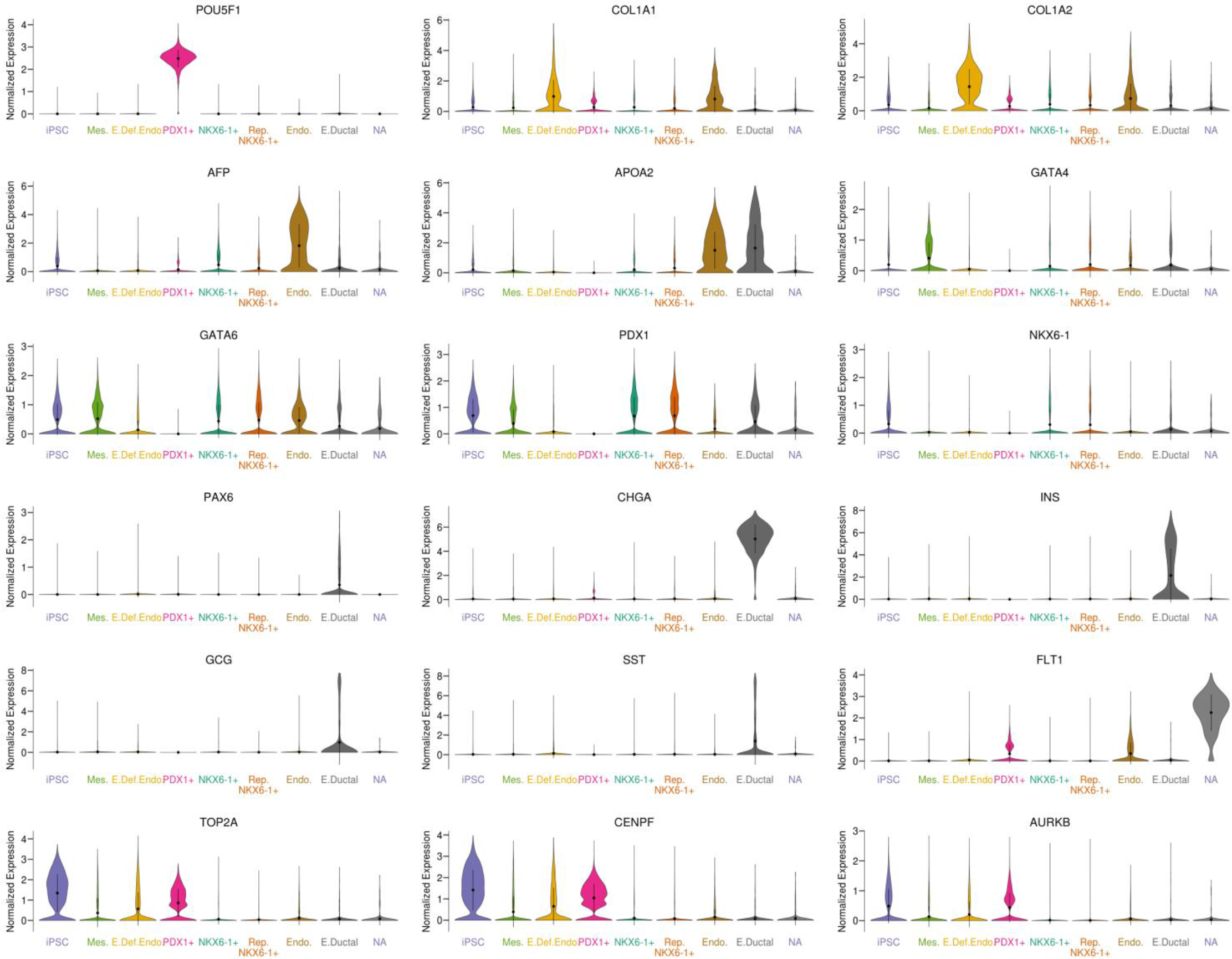
Expression of 17 marker genes in each of the eight scRNA-seq clusters. Violin plots showing the distributions of normalized expression for marker genes described in Figure 1F for each scRNA-seq cluster in Figure 1B.

**Figure S6:**
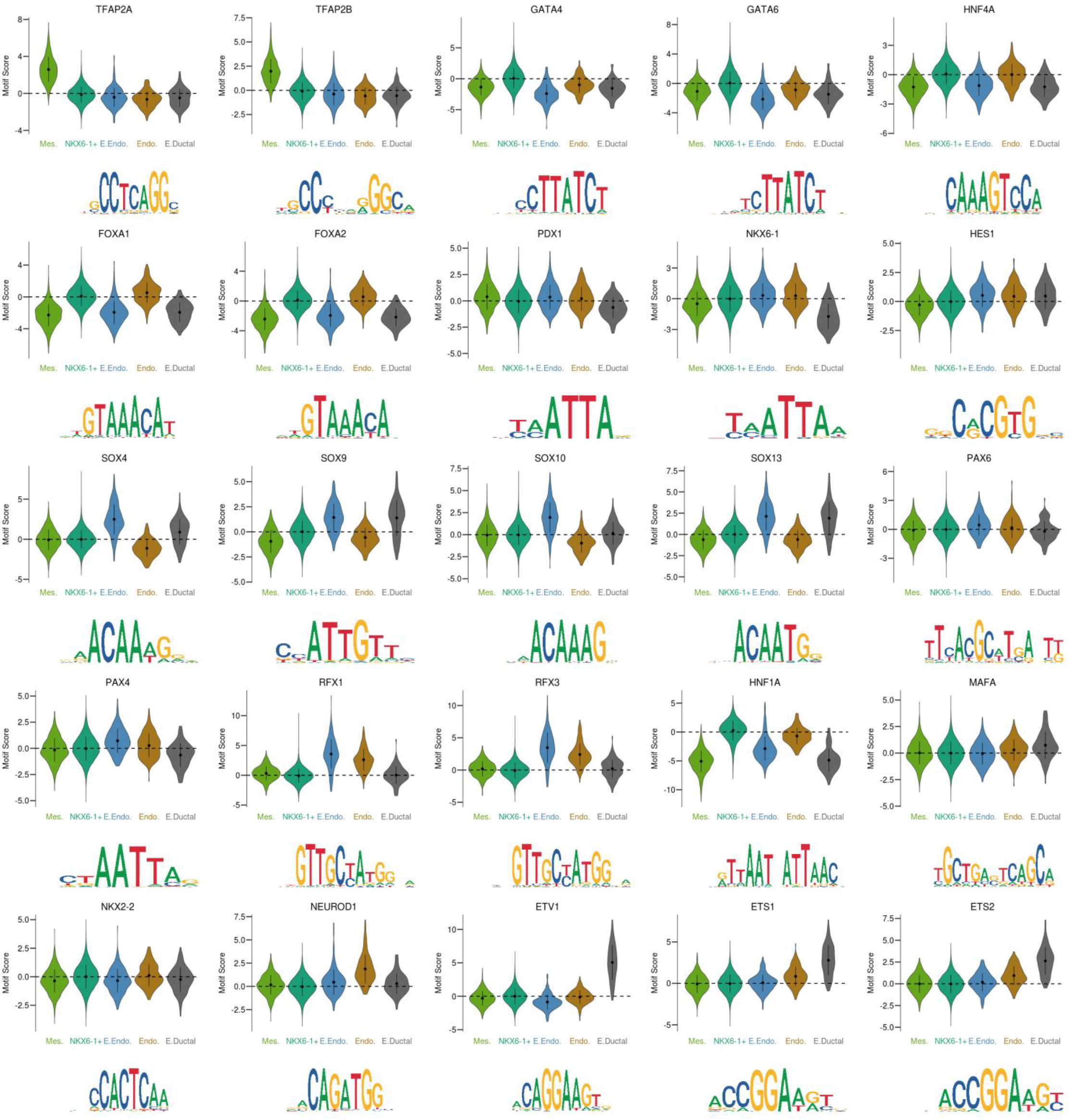
Motif activity of 23 transcription factors in each of the five snATAC-seq clusters. Violin plots showing the distribution of chromVAR motif activity score for the transcription factors described in Figure 1G for each snATAC-seq clusters in Figure 1D. Motif logos are shown underneath the corresponding plot.

**Figure S7:**
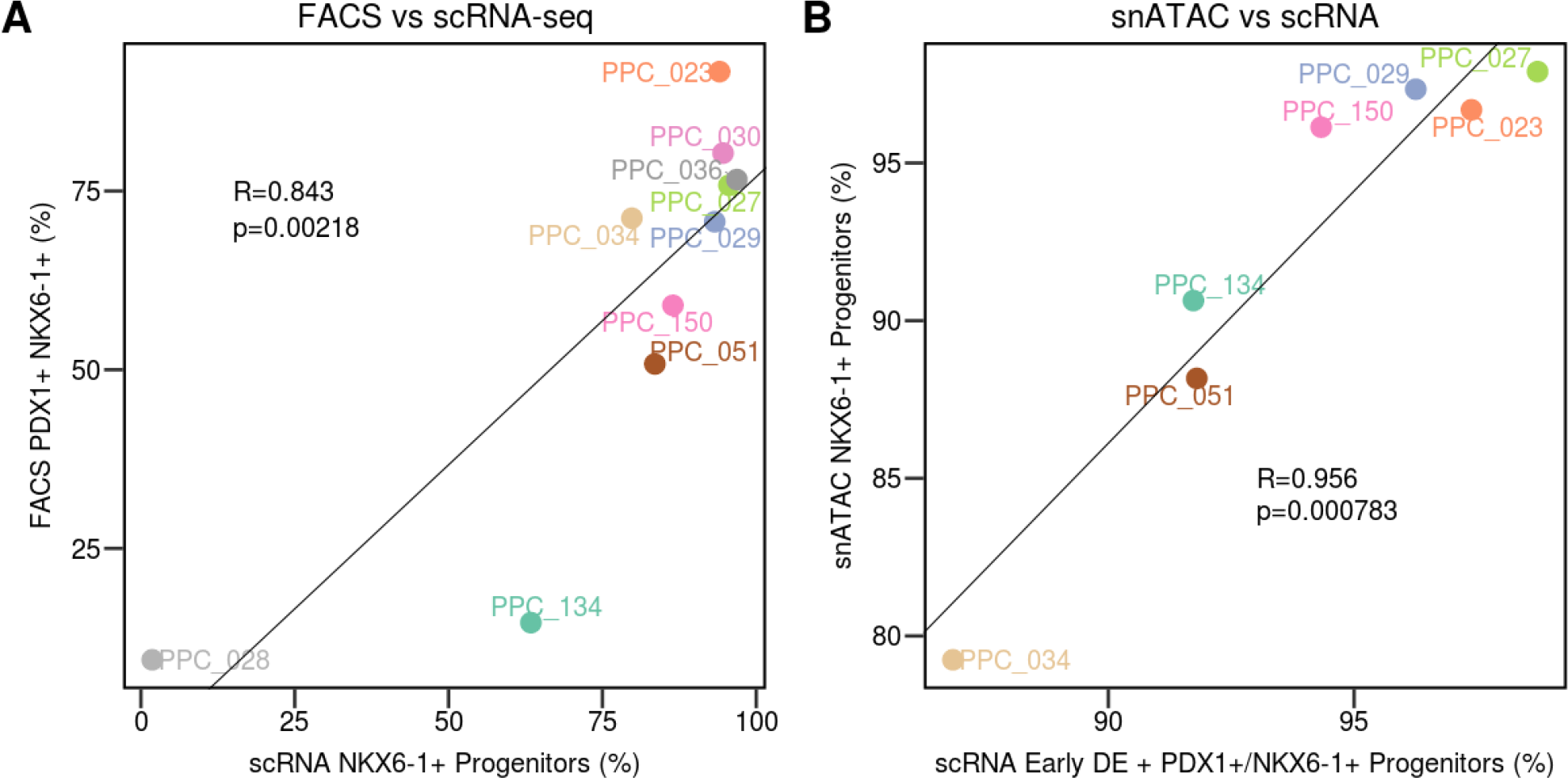
Correlation between flow cytometry, scRNA-seq and snATAC-seq results. To determine the correspondence between FACS, scRNA-seq, and snATAC-seq, we compared the fraction of cells expressing both *PDX1* and *NKX6-1* within each iPSC-PPC sample. For scRNA-seq, we computed the total number of NKX6-1+ progenitors as the sum of NKX6-1+ progenitors and replicating NKX6-1+ progenitors (X axis, Figure S7A), as these cells express both *PDX1* and *NKX6-1*. We found that the FACS and scRNA-seq was significantly correlated (R = 0.843, p = 0.00218, Figure S7A). We next determined whether the cell type fractions in snATAC-seq corresponds to scRNA-seq. Because snATAC-seq was not able to detect PDX1+ progenitors and early definitive endoderm (DE), we reasoned that these cells may be included in the main NKX6-1+ progenitor nuclei cluster. Therefore, we computed the fraction of cells that are early DE, PDX1+, NKX6-1+ progenitors and replicating NKX6-1+ progenitors in scRNA-seq and compared it to the fraction of nuclei that are NKX6-1+ progenitors in snATAC-seq. We found a significant correlation between scRNA-seq and snATAC-seq (R = 0.956, p = 0.000783, Figure S7B). These results show that scRNA-seq, snATAC-seq, and FACs are highly associated with each other, and that both sequencing methods can capture the variable differentiation efficiency in iPSC-PPC.

**Figure S8:**
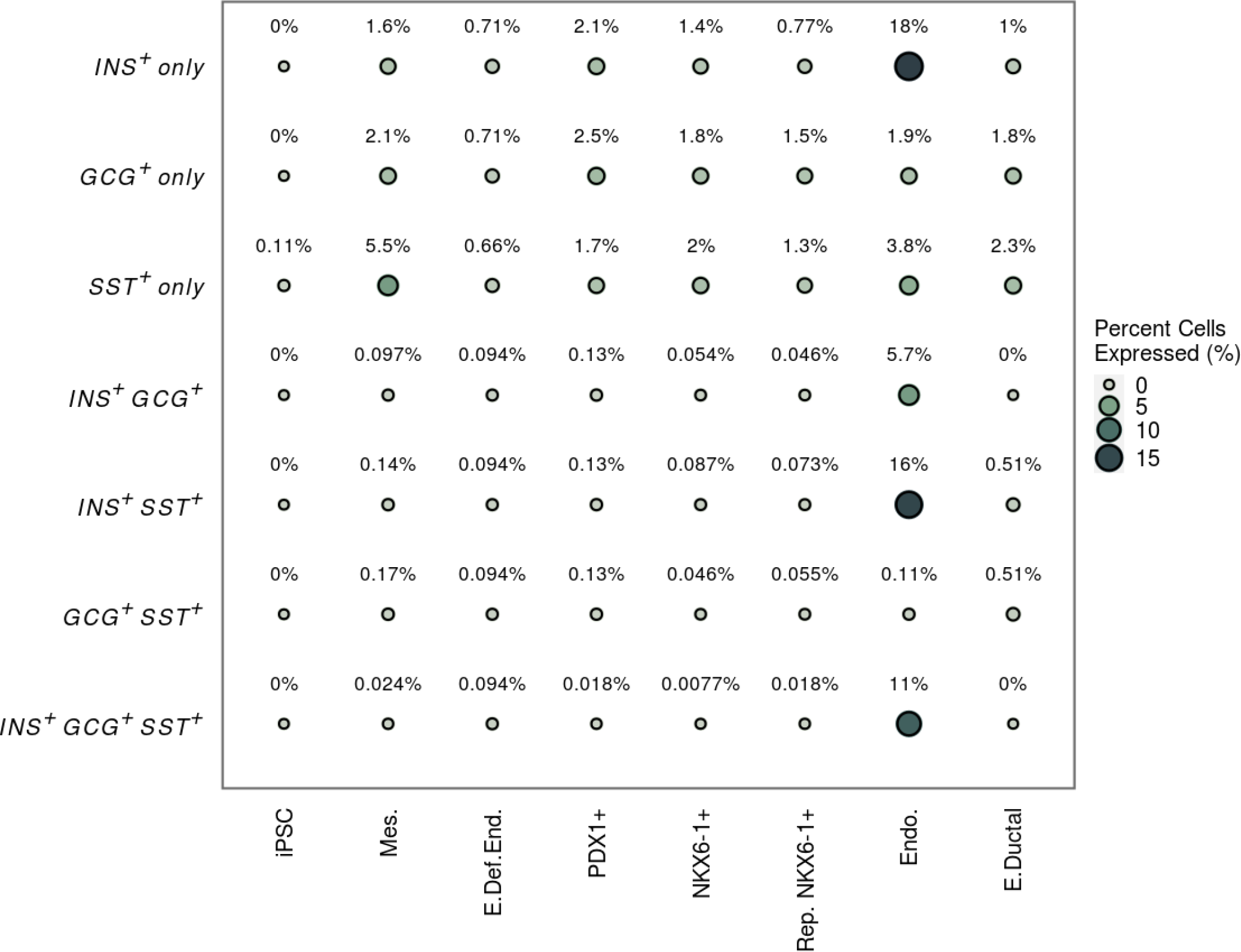
Co-expression of insulin, glucagon, and somatostatin in pancreatic progenitors. Bubble plot showing the percentage of cells that express endocrine-specific hormones in scRNA-seq clusters, where radius and shade indicate the percentage of cells expressing above 10% of the maximal expression for the indicated gene across all cells.

**Figure S9:**
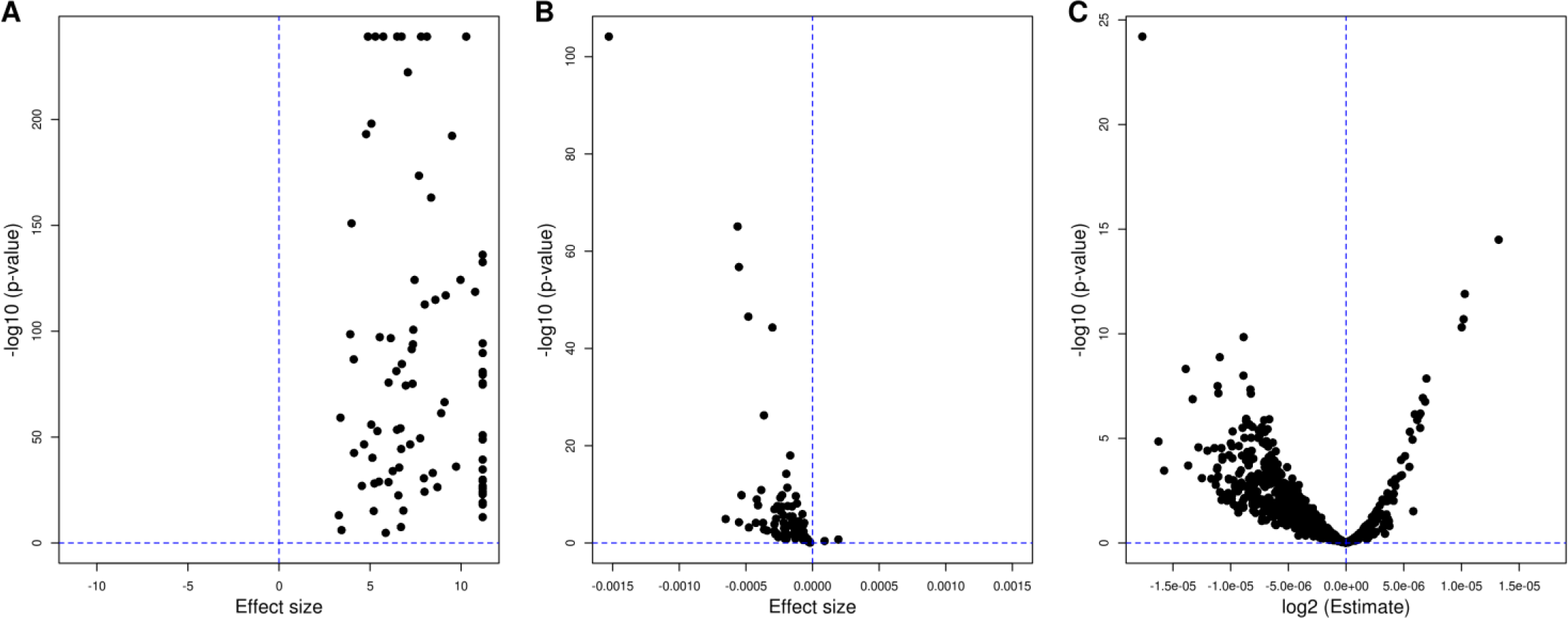
Associations between deltaSVM scores, ASE and transcription factor footprints. Volcano plots showing the associations between deltaSVM scores, ASE and transcription factor footprints. Each point represents a transcription factor. (A) For each of the 89 transcription factors tested with both deltaSVM and TOBIAS, we investigated the agreement between these two tools. Here, we determined if variants predicted to have allelic effects on a specific transcription factor by deltaSVM were more likely than expected to occur at genomic locations bound to the same transcription factor, as determined by TOBIAS. Effect size (X axis, log2 ratio) and p-values (Y axis, Fisher’s exact tests) were measured using the *fisher.test* function in R. All tests had significant p-values after FDR correction (Benjamini-Hochberg) and had a positive log2 ratio and, overall, the distribution of log2 ratios was significantly greater than zero (p = 2.2 x 10^-47^, t-test, measured with the *t.test* function in R, with option *mu = 0*). Each dot represents one of the 89 tested transcription factors. (B) For the same transcription factors in (A), we tested if variants predicted to have allelic effects by deltaSVM were more likely to be closer to transcription factor footprints determined using TOBIAS than expected. We computed the distance of each variant from the closest transcription factor footprint on the same peak, and performed a linear regression between the distance and the absolute value of the variant score measured by deltaSVM. For all but two transcription factors, we observed a negative association (X axis = effect size; Y axis = p-value, measured using the *lm* function in R) and, overall, the effect size distribution was significantly lower than zero (p = 4.3 x 10^-15^, t-test, measured with the *t.test* function in R, with option *mu = 0*), indicating that variants closer to transcription factor footprints are more likely to have stronger allelic effects. (C) We tested the association between the major allelic frequency of each heterozygous variant in iPSC-PPC peaks tested for ASE and the distance from its closest footprint for each of the 746 transcription factors analyzed using TOBIAS. For most transcription factors, we observed a negative association (X axis = effect size; Y axis = p-value, measured using the *lm* function in R) and, overall, the effect size distribution was significantly lower than zero (p = 6.45 x 10^-84^, t-test, measured with the *t.test* function in R, with option *mu = 0*), indicating that variants closer to transcription factor footprints are more likely to have stronger allelic effects.

**Figure S10:**
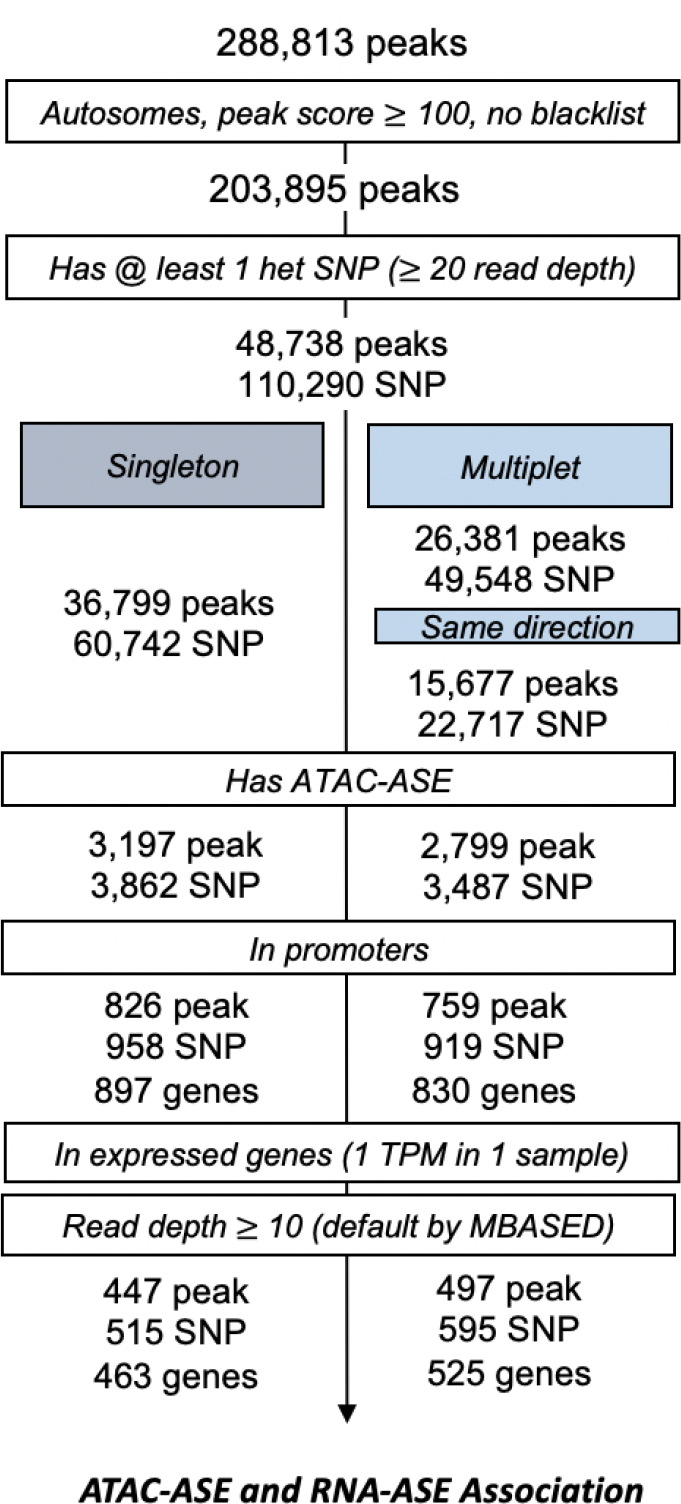
Workflow for computing allele-specific effects (ASE) in snATAC-seq peaks. Peaks on autosomal chromosomes were filtered based on: peak score ≥ 100, not within ENCODE blacklist regions, and if they contained at least one heterozygous SNP in at least one of the seven snATAC-seq samples. SNPs were further categorized as singletons (i.e. observed in only one individual) or multiplets (i.e. observed in more than one individual). Multiplets with consistent allele direction in all individuals were retained. SNPs were then tested for ASE using a two-sided binomial test assuming a null hypothesis that both alleles were observed at equal proportions. We classified SNPs as having ASE if FDR ≤ 0.05 and major allele frequency ≥ 0.6. SNPs with ASE at promoter regions were further examined for ASE association with gene expression using bulk RNA-seq (Figure 2D).

## TABLE LEGENDS

Table S1: Study information, including subject details, differentiation efficiency, and generated molecular data from 1 iPSC and 10 iPSC-PPCs

In Sheet 1: Subject_UUID is the assigned Universal Unique Identifier (UUID) for each subject (Column A) used in this study. Sex (Column B), age (Column C), and ethnicity (most similar 1KG population; Column D) are provided. iPSCORE_Family (Column E) are the family identifiers used to identify related family members. Columns A-E are shown as included in dbGaP (phs001325.v1.p1; phs000924.v1.p1) as part of the iPSCORE Resource.

In Sheet 2: We provide information for each sample in our study. Subject_UUID is provided in Column A. Cell_type (Column B) indicates the type of cell (iPSC or PPC) obtained from each subject as part of this study. Unique Differentiation Identifier (UDID, Column C) is a unique digit assigned for each attempted iPSC-PPC differentiation. %PDX1+ (Column D), %NKX6.1+, (Column E), and %PDX1+_NKX6.1+ (Column F) are the fractions of cells from each iPSC-PPC differentiation positively stained for PDX1, NKX6-1, or both PDX1 and NKX6-1, respectively. Data type indicates the type of sequencing method performed for each differentiation (Column G). The UUID for each sequenced sample is provided (Column H). Pooling schemes for samples combined prior to sequencing are shown (Column I).

In Sheet 3: We provide the UUIDs of each iPSC-PPC sample (Column A) by subject (Column B), WGS (Column C), bulk RNA-seq (Column D), fresh scRNA-seq (Column E), cryopreserved scRNA-seq (Column F), and snATAC-seq (Column G). Bulk RNA-seq was generated for all the of 10 iPSC-PPC samples that have scRNA-seq, but in this study we analyzed only the seven that have matched snATAC-seq (Table S1).

Table S2: scRNA-seq metadata

The table shows, for each of the 83,971 single cells, the sample UDID (Column A), the preparation of the sample corresponding to either fresh or cryopreserved (Column B), the cell barcode (Column C), UUIDs for subject, WGS, and scRNA-seq (Columns D-F), the number of reads (Column G), the number of genes detected (Column H), the percent of mitochondrial reads (Column I), the cell type assignment (Column J), UMAP coordinates (Column K-L), and the cluster assignments at resolutions 0.05, 0.08, and 0.1 (Columns M-O). The UUIDs for WGS was obtained using Demuxlet (Kang et al., 2018) and then mapped to subject and sample UUID. Barcodes for cells from freshly prepared samples were formatted as *barcode*-*aggregate_id* (Sheet 2) while those from cryopreserved samples were formatted as *barcode-1*.

Because this table’s size is too large, it has been deposited on figshare: https://doi.org/10.6084/m9.figshare.15109581

A Seurat R object including all scRNA-seq data has been deposited on figshare: https://doi.org/10.6084/m9.figshare.15109422

Table S3: Genes differentially expressed in scRNA-seq clusters

For each gene (Column A) and each scRNA-seq cluster (Column B), we computed Wilcoxon rank sum test between normalized expression values across cells within the cluster and cells outside of the cluster. The table provides the average log2 fold-change between the groups (Column C), the average expression for each group (Column D-E), the fraction of cells with expression greater than 0.1 (Column F-G), the p-value (Column H), and q-value adjusted by Bonferroni correction (Column I). Genes with q-value ≤ 0.05 were considered differentially expressed.

Table S4: snATAC-seq metadata

The table shows, for each of the 26,564 single nuclei, the cell barcode (Column A), the sample UDID (Column B), UUIDs for subject, WGS, snATAC-seq, and matched scRNA-seq (Columns C-F), quality control parameters from CellRanger-ATAC and Signac (Columns G-V), the assigned cell type (Column W), UMAP coordinates (Column X-Y), and the cluster assignments at resolutions 0.1, 0.15, and 0.2 (Columns Z-AB). The UUIDs for WGS were obtained using Demuxlet (Kang et al., 2018) and then mapped to subject and sample UUID (snATAC-seq and scRNA-seq). Because this table’s size is too large, it has been deposited on figshare: https://doi.org/10.6084/m9.figshare.15109581 A Seurat R object including all snATAC-seq data has been deposited on figshare: https://doi.org/10.6084/m9.figshare.15109422

Table S5: Peaks differentially expressed in snATAC-seq

For each cluster in snATAC-seq described in Figure 1D, we performed differential expression using the *FindAllMarkers* function in Signac. The table provides the cluster name, peak ID, average log2 fold-change, the fraction of cells within cluster that has peak expression > 0.1, the fraction of cells outside of cluster that has peak expression > 0.1, p-value, and q-value adjusted with a Bonferroni correction. We consider peaks with q-value ≤ 0.05 to be differentially expressed.

Table S6: Motifs enriched in snATAC-seq peaks

We performed motif enrichment analyses for 633 transcription factors in the JASPAR 2020 database. The table shows, for each snATAC-seq cluster, the tested motif ID and name from JASPAR, the p-value from Wilcoxon rank sum test, and the adjusted p-value using a Bonferroni correction. Motifs with q-value ≤ 0.05 were considered differentially enriched.

Table S7: Evidence of SNPs for association with transcription factor binding or allelic effects

In total, we were able to investigate the functional associations for 349,572 variants, including 325,942 common SNPs (allele frequency > 5%) in the iPSCORE collection and 110,290 heterozygous variants in the seven iPSC- PPC samples (86,660 variants were in common). The table shows, for each variant, its associated peak, whether it belongs to the 325,942 common SNPs or to the 110,290 heterozygous variants and its functional associations: 1) overlap with transcription factor footprints (TOBIAS); 2) prediction of allele-specific effects (deltaSVM); or 3) ASE in snATAC-seq. Because this table’s size is too large, it has been deposited on figshare: https://doi.org/10.6084/m9.figshare.15109581 R objects including all results from TOBIAS, including the position of all bound transcription factor binding sites and all the overlaps between SNPs and transcription factor binding sites, have been deposited on figshare: A Seurat R object including all scRNA-seq data has been deposited on figshare: https://doi.org/10.6084/m9.figshare.15109422

Table S8: deltaSVM prediction of allele-specific transcription factor binding

The table shows all associations between each of the 325,942 SNPs in snATAC-seq peaks and each of the 94 tested transcription factors. For each SNP, shown are: its chromosome and position, reference and alternative alleles, tested transcription factor, oligo binding score for reference and alternative allele (Yan et al., 2021), whether deltaSVM predicts that the transcription is bound, deltaSVM score for allelic TF binding, and the consequence of the SNP on the transcription factor binding (“Gain”: the alternative allele has a stronger binding score; “Loss”: the reference allele has a stronger binding score; “None”: no difference between reference and alternative alleles). Only the 52,653 SNPs predicted to be associated with allele-specific transcription factor binding are shown. A table including all 30,638,548 tested SNPs/transcription factor pairs has been deposited on Figshare (https://doi.org/10.6084/m9.figshare.15109581).

Table S9: Associations between transcription factor binding site predictions from deltaSVM and TOBIAS

The table shows the enrichment for transcription factor binding and allelic effects predicted by deltaSVM in the genomic regions associated with a bound transcription factor annotated by TOBIAS. For each of the 89 transcription factors (the transcription factor ID by JASPAR and the transcription factor name by deltaSVM are shown in the table) in common between deltaSVM predictions and TOBIAS, we performed two enrichment analyses (shown in Figure S9A,B): 1) we tested for the enrichment of variants predicted to overlap a transcription factor binding site by deltaSVM and bound transcription factor binding sites measured by TOBIAS, using Fisher’s exact test (the table shows estimate, p-value and q-value adjusted using Benjamini-Hochberg’s method); and 2) we tested for the enrichment of variants predicted by deltaSVM to have allelic effects at a transcription factor binding site and bound transcription factor sites measured by TOBIAS, using linear regression (the table shows effect size, standard error, p-value and q-value adjusted using Benjamini-Hochberg’s method).

Table S10: Allele-specific effects of heterozygous SNPs in snATAC-seq peaks

Shown are the allele-specific effects of heterozygous SNPs with read depth ≥ 20 in heterozygous iPSCORE individuals (Column A). Variant ID indicates the chromosome position, reference allele, and alternate allele of the heterozygous SNP (Column B). The peak that overlaps with the SNP (Column C), the peak score from MACS2 (Column D) and the gene ID and name whose promoter overlaps with the SNP (Column E-F) are provided. The number of reads calculated by *samtools mpileup* from snATAC-seq BAM files overlapping the reference allele, alternative allele, or both are shown (Columns G-I). The major allele frequency was calculated as the fraction of reads that map to the allele with greater number of reads (Column J). Information about whether the variant is a singleton (i.e. observed only in one individual) (Column K) or observed in a single allele direction are given (Column K-L). P-values (Column M) were calculated using binomial test with the alternative hypothesis that both alleles were observed in equal frequency. P-values were adjusted using Benjamini-Hochberg’s method (Column N). SNPs showed allele-specific effects if FDR ≤ 0.05 and the major allele frequency is ≥ 0.6 (Column O).

Because this table’s size is too large, it has been deposited on figshare: https://doi.org/10.6084/m9.figshare.15109581

Table S11: deltaSVM prediction of allele-specific transcription factor binding

The table shows the associations between deltaSVM scores and the alternative allele frequency of all SNPs overlapping bound transcription factors. Shown are: the SNP ID (as in Table S4), transcription factor, deltaSVM score and allele frequency calculated from snATAC-seq.

Because this table’s size is too large, it has been deposited on figshare: https://doi.org/10.6084/m9.figshare.15109581.

Table S12: Associations between transcription factor binding and ASE

The table shows the associations between the major allele frequency of variant tested for ASE and its distance from each bound transcription factor binding site in the same snATAC-seq peak measured by TOBIAS (Figure S9C). The table shows, for each of the tested 746 transcription factors, its JASPAR ID, the measured effect size, standard error, p-value and q-value adjusted using Benjamini-Hochberg’s method.

Table S13: Correspondence between ASE at promoters and their associated genes

Shown are the allele-specific effects of 5,380 expressed genes that have a snATAC-seq peak at the promoter region. The analysis was performed using MBASED (Mayba et al., 2014) on a per-sample basis. For each sample and gene (Columns A-C), we provide information about the major allele frequency (Column D), p-value of heterogeneity (Column E), p-value of ASE (Column F), FDR-corrected p-value using Benjamini Hochberg’s method (Column G), and whether this gene exhibits ASE or not (Column H). We determine a gene to exhibit ASE if FDR ≤ 0.05 and the major allele frequency is ≥ 0.6. The major allele frequency, p-value of ASE, and p-value of heterogeneity were computed by MBASED using the 1-sample analysis in the provided protocol.

Table S14: Associations between T2D-associated SNPs and allelic effects in iPSC-PPC

For each of the 66,600 SNPs in T2D credible sets (Mahajan et al., 2018), the table shows: their associated locus, including the position of the index SNP, the index gene and the RS ID of the index SNP, as defined in Mahajan et al. (Mahajan et al., 2018); the PPA and the rank in the credible set; whether each SNP overlaps an iPSC-PPC snATAC-seq peak; whether it is predicted to have allelic effects, based on deltaSVM results; and whether it has ASE in the iPSC-PPC snATAC-seq dataset.

